# Generating parallel representations of position and identity in the olfactory system

**DOI:** 10.1101/2022.05.13.491877

**Authors:** István Taisz, Erika Donà, Daniel Münch, Shanice N. Bailey, Billy J. Morris, Kimberly I. Meechan, Katie M. Stevens, Irene Varela, Marina Gkantia, Philipp Schlegel, Carlos Ribeiro, Gregory S.X.E. Jefferis, Dana S. Galili

**Affiliations:** Neurobiology Division, MRC Laboratory of Molecular Biology, United Kingdom; Champalimaud Foundation, Lisbon, Portugal; Department of Zoology, University of Cambridge, United Kingdom

**Keywords:** social behavior, sexual dimorphism, sensory physiology, pheromones, connectomics, neural circuits

## Abstract

Sex pheromones are key social signals in most animals. In *Drosophila* a dedicated olfactory channel senses a male pheromone, cis-vaccenyl acetate (cVA) that promotes female courtship while repelling males. Here we show that flies use separate cVA processing streams to extract qualitative and positional information. cVA olfactory neurons are sensitive to concentration differences in a 5 mm range around a male. Second-order projection neurons detect inter-antennal differences in cVA concentration, encoding the angular position of a male. We identify a circuit mechanism increasing left-right contrast through an interneuron which provides contralateral inhibition. At the third layer of the circuit we identify neurons with distinct response properties and sensory integration motifs. One population is selectively tuned to an approaching male with speed-dependent responses. A second population responds tonically to a male’s presence and controls female mating decisions. A third population integrates a male taste cue with cVA; only a simultaneous presentation of both signals promotes female mating via this pathway. Thus the olfactory system generates a range of complex percepts in discrete populations of central neurons that allow the expression of appropriate behaviors depending on context. Such separation of olfactory features resembles the mammalian what and where visual streams.

**Highlights:**

- cVA male pheromone has a 5 mm signaling range, activating two parallel central pathways
- Pheromone-sensing neurons have spatial receptive fields sharpened by contralateral inhibition
- Position (where) and identity (what) are separated at the 3rd layer of cVA processing
- Integrating taste and cVA in sexually dimorphic aSP-g controls female receptivity

## Introduction

Olfaction allows animals to identify and evaluate objects, but also to gather spatial information about the environment (Baker et al., 2018; von Békésy, 1964). Studies of primate visual cortex show that object identity and motion are processed in parallel ventral and dorsal streams, also called the *what* and *where* pathways (Goodale and Milner, 1992). This separation follows the logic of invariant representation: recognizing the identity of an object is largely independent of its motion, while object motion is independent of identity (Bakhtiari et al., 2021; Cadieu and Olshausen, 2012). When a neuron represents object identity, it is active as long as the object is present; a tonic response. Motion on the other hand must be extracted from time-varying signal correlations generating phasic neuronal responses. These separate processing strategies have been extensively studied in vision, but a comprehensive, synaptic resolution understanding of the circuit mechanisms remains a major challenge.

Olfaction is an anatomically shallow sensory system. In mammals and invertebrates just two synapses separate the sensory periphery from neurons that store memories or organize innate behaviors (Dolan et al., 2018; Frechter et al., 2019; Knaden and Hansson, 2014; Li and Liberles, 2015; Owald and Waddell, 2015; Wilson and Mainen, 2006). Olfactory cues are key signals for social interactions in most animals. For example in *Drosophila*, males produce cis-vaccenyl acetate (cVA), a low-volatility pheromone which acts as a female aphrodisiac but promotes aggression in males (Kurtovic et al., 2007; Wang and Anderson, 2010). These genetically-specified, sexually dimorphic behaviors provide a robust read-out for the qualitative information (e.g. sexual and species identity) carried by this pheromone. The prototypical pheromone, bombykol, is remarkable in its ability to guide male moths over kilometers, simultaneously conveying both positional and identity information. However most social behaviors occur at closer range and are multimodal. Both male and female flies use visual, olfactory, and gustatory cues to identify conspecifics and organize sexually dimorphic behaviors including courtship and aggression (Cook, 1979; Rings and Goodwin, 2019; Spieth, 1952; Wang et al., 2011). In forming a binary decision about whether to accept a potential mate, females integrate cues like cVA, courtship song, and male contact pheromones (Bussell et al., 2014; Crossley et al., 1995; Grillet et al., 2006; Spieth, 1952). A sexually dimorphic neuron cluster, pC1, and its male homologue P1, have been identified as a central brain hub for sexual behaviors (Hoopfer et al., 2015; Rezával et al., 2016; Wang et al., 2020b; Yamamoto et al., 2008; Zhou et al., 2014), with clear analogies to circuit modules in the mammalian hypothalamus (Anderson, 2016).

Sex pheromones have proven a powerful entry point to study the genetic and circuit basis of behavior (Auer and Benton, 2016; Li and Dulac, 2018). However, there remain large gaps in our understanding, even in favorable models like *Drosophila* and its canonical pheromone, cVA. cVA is synthesized internally within the male and passed to the female during mating (Brieger and Butterworth, 1970) but it is currently unclear when and where it acts during social behavior: does it act as a diffuse permissive signal or do stimulus location and strength convey important information? If so, how can these be detected? Within the brain, a second-order interneuron has been identified that receives cVA information (Datta et al., 2008), but puzzlingly, manipulations have not linked neuronal activity to female receptivity. At the third-order, two populations of cVA-responsive interneurons have been identified (Cachero et al., 2010; Ruta et al., 2010) and we have previously shown that they form a sexually dimorphic circuit switch (Kohl et al., 2013). Nevertheless the behavioral significance of these neurons in courtship remains untested. Finally pC1 neurons are cVA-responsive (Zhou et al., 2014), yet their cVA inputs have not been found.

Here we provide a systems level structural, physiological and behavioral characterization of three layers in the cVA processing circuit. We use connectomics to find novel second- and third-order neurons, revealing an unexpectedly concise pathway from sensory neurons to pC1 central integrators. We find that male flies are surrounded by a narrow pheromone halo; presenting a fly mounted on a micromanipulator shows that olfactory neurons have sub-millimeter precision spatial receptive fields. Comparing pheromone signals from both antennae – using an active contrast circuit that we identify through the interplay of connectomics and physiology – effectively allows flies to “see” each other in the dark using smell. Parallel and hierarchical processing generates a wealth of sensory percepts including features of both position and identity. These different circuit elements enable a sex pheromone to mediate both sexually dimorphic and shared behaviors. Our findings substantially advance understanding of an influential model system, describing a complete sensory processing hierarchy in unprecedented detail. More generally, they show that olfaction has surprisingly strong analogies with other sensory systems. Like the auditory system, positional information is synthesized from active comparison of bilateral sensory signals, while separation of *what* and *where* pathways is reminiscent of deeper layers of visual cortex.

## Results

### Parallel cVA pathways have distinct effects on sexual behaviors

EM (electron microscopy) connectomics is presently unique in revealing complete circuits with synaptic resolution; this is particularly powerful when applied at whole brain scale, for which adult *Drosophila* represents the current state of the art. Through tracing, annotation and analysis of two EM datasets, we have been able to obtain a comprehensive structural framework to understand neural information processing of the pheromone cVA. We began by tracing downstream partners of cVA responsive olfactory receptor neurons (ORNs), which express receptor Or67d and target the DA1 glomerulus (Figure 1A) using the Full Adult Fly Brain (FAFB) dataset (Bates et al., 2020a; Zheng et al., 2018). In addition to the well-known uniglomerular DA1 lateral PNs (lPNs) and inhibitory ventral PNs (vPNs; Figure 1B, Figure S1A, I, (Datta et al., 2008; Marin et al., 2002; Wong et al., 2002), we found a novel uniglomerular cell type from the lateroventral lineage, which we therefore called DA1 lvPN; we later identified the same cell type in the hemibrain EM volume (M_lvPNm45 (Scheffer et al., 2020; Schlegel et al., 2021). DA1 lvPNs receive 99% of their sensory input from Or67d ORNs and make the same axonal projections in both sexes; like lPNs they relay cVA information to the lateral horn (LH) but significantly they bypass the mushroom body associative learning center, instead projecting to the superior intermediate protocerebrum (SIP), a multimodal higher-order neuropil (Figure 1C, Figure S1C). We used the EM morphology to obtain a split GAL4 driver line (Figure 1C, Figure S1B) and confirmed this is a cholinergic, excitatory cell type (Figure S1E); *in vivo* 2-photon calcium imaging showed robust cVA responses (Figure 1D). lPNs and lvPNs therefore form parallel excitatory cVA processing pathways.

**Fig. 1.**
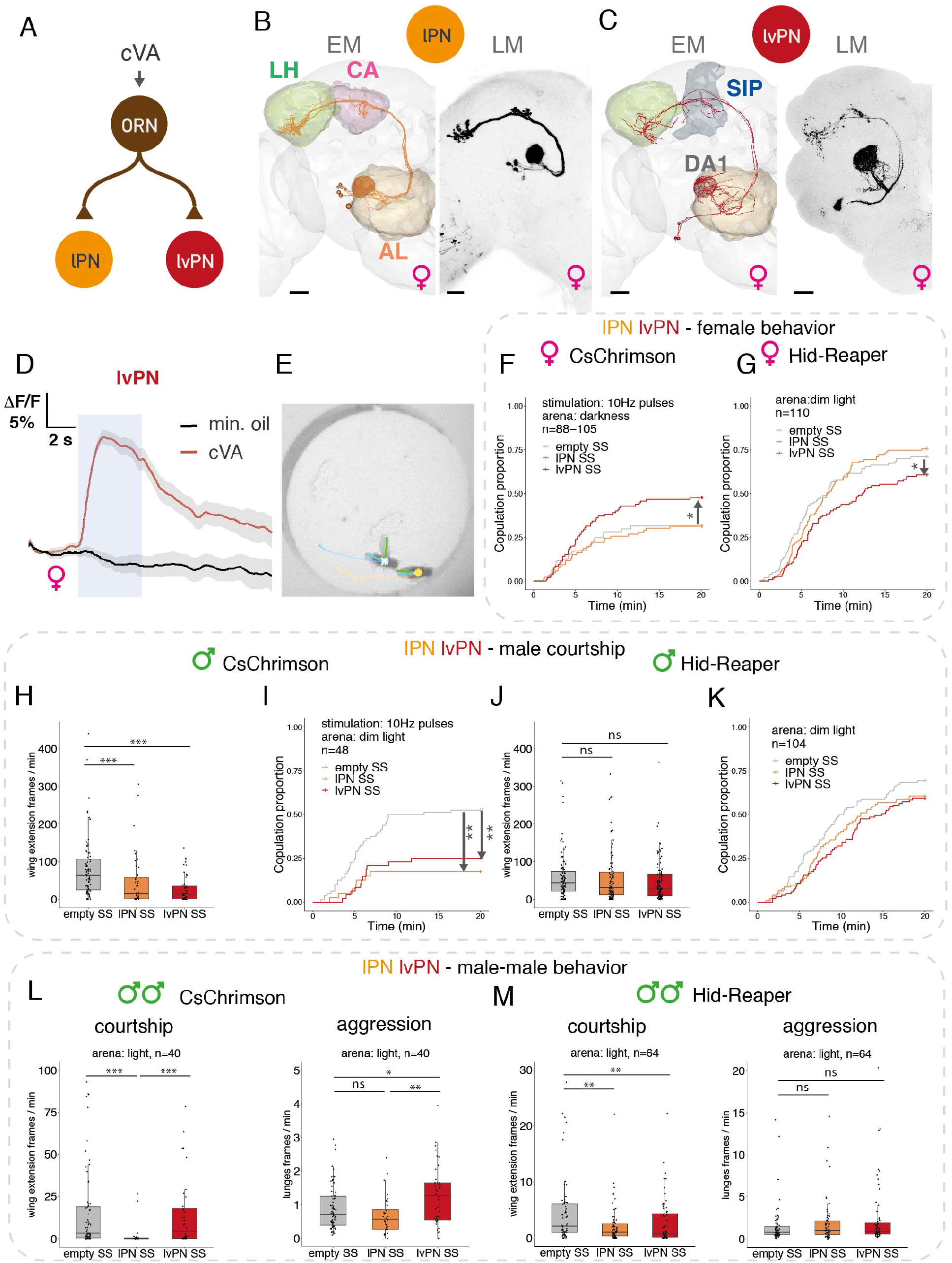
Parallel cVA pathways have distinct effects on sexual behaviors. **A**: Connectivity of Or67d ORNs, DA1 IPN, and DA1 IvPNs based on the hemibrain. Number of synaptic connections: ORN - IPN: 9187 across 7 IPNs; ORN - lvPN: 286 across 3 lvPNs. **B, C**: EM reconstructions in FAFB (left) and light microscopy (LM) images (right) of DA1 lPNs (B) and DA1 lvPNs (C). Maximum intensity projections of reporter expression driven in female brains by lPN stable split line (SS) or lvPN SS, respectively, are shown. (AL: antennal lobe, CA: calyx, LH: lateral horn, SIP: superior intermediate protocerebrum, DA1: dorsal anterior 1 glomerulus). Scale bars: 20 μm. **D**: cVA activates lvPNs. GCaMP6s responses in lvPN axons to cVA presentation (10%) and solvent control. Shaded blue area: odor delivery (5 s). Average response from 6 flies, 6 presentations, gray area is the standard error of the mean (SEM). **E**: Image of a chamber used for courtship essay (see methods). Fly centroids, wing positions, and fly trajectories are annotated via FlyTracker and plotted. Scale bar: 2 mm. **F, G:** Manipulating DA1 lvPN SS and lPN SS in virgin females paired with wildtype males. (F) Optogenetic activation of DA1 lvPNs increased female receptivity, while activating lPNs had no effect. (G) Hid, Reaper-induced ablation of lvPNs decreased female receptivity, while ablating lPNs had no effect. **H, I, J, K:** Manipulating DA1 lPN SS or lvPN SS in males paired with wildtype females. Optogenetic activation decreased courtship (H) and mating proportion (I). Hid,Reaper-induced ablation had no effect on male courtship (J) or mating proportion (K). **L, M:** Manipulating lPN SS or lvPN SS in pairs of males. (L - left) Optogenetic activation of lvPN increased male-male aggression, while activating lPN had no effect. (L - right) lPN activation reduced male-male wingextension, while activating lvPN had no effect. Hid,Reaper-induced ablation had no effect on aggression (M - left) and reduced wing extension (M - right). Throughout the figures: mating curves represent the proportion of mated females over time. ‘Pulses’= 10Hz red light pulses given for 5s on - 5s off, while ‘constant’= constant red light on; 627 nm, 8 μW/mm^2^ during 20 min recording, in an otherwise complete dark incubator. * p<0.05; ** p<0.01; *** p<0.001. See genotypes and statistics in Supplementary File 1.

How do these parallel PN pathways contribute to the sexspecific effects of cVA on courtship and aggression? We measured sexual behaviors in pairs of virgin flies freely interacting for 20 minutes (Figure 1E), while optogenetically activating each PN type by CsChrimson (Klapoetke et al., 2014). We tracked fly and wing position using FlyTracker (Eyjólfsdóttir, 2014), measured copulation latency and trained classifiers for discrete sexual behaviors using JAABA (Kabra et al., 2013). Activating lPNs in virgin females paired with wildtype males had no effect on mating. However, lvPN activation in females increased copulation rate, reflecting higher female receptivity compared to controls (Figure 1F). Cell killing experiments had consistent effects: lvPN ablation reduced mating success while lPN ablation had no effect (Figure 1G).

Turning our attention to males, optogenetic activation of either lPNs or lvPNs resulted in decreased courtship towards wildtype females and strongly reduced copulation rate (Figure 1H, I); the behavioral significance of both PN types contrasts with the female results earlier and likely reflects sex differences in downstream architecture. Since only males produce cVA and transfer it to females upon mating, it is not surprising that genetic ablation had no effect on male courtship towards virgin females (Figure 1J, K). Males regularly court other males even though this is actively suppressed by cVA and contact pheromones (Billeter et al., 2009). We found that male-male courtship was strongly reduced by lPN (but not lvPN) activation (Figure 1L); ablating lPNs reduced male-male courtship with a weaker effect for lvPNs (Figure 1M). Another prominent role of cVA in males is promoting aggression. We found that lvPN, but not lPN, activation, moderately increased aggression between male pairs assayed in the same arena (Figure 1L). Ablating lPNs or lvPNs had no effect on male-male aggression (Figure 1M), likely due to low baseline levels of aggression in group-reared flies. Finally, we saw no effect on female-female aggression by activating either lPNs or lvPNs in pairs of females (Figure S1D, G).

In summary, the newly characterized lvPNs promote many of the behavioral effects of cVA, increasing female receptivity and malemale aggression. lPN manipulations recapitulated only the courtship suppressing effects of cVA in male flies and had no effect on female sexual behavior. This explains why until now attempts to link cVA processing to behavior have largely stalled in the sensory periphery.

### Parallel cVA pathways differentially signal male distance and sustained presence

Why are there two PN pathways with different behavioral effects? Both receive cVA sensory input, but they could still extract distinct stimulus features. We carried out detailed sensory physiology using a male fly as a stimulus (Clowney et al., 2015; Tachibana et al., 2015) to ensure that we were providing ethologically relevant odor concentrations. Crucially, we mounted the male on a micromanipulator so that we could present it at precisely defined positions relative to the observer fly, and measured calcium signals in DA1 ORN, lPN, or lvPN axons (Figure 2A inset, Figure S2A, B). This provided much more precise control of cVA concentration at the antenna while also mimicking the sensory experience of flies interacting with each other.

**Fig. 2.**
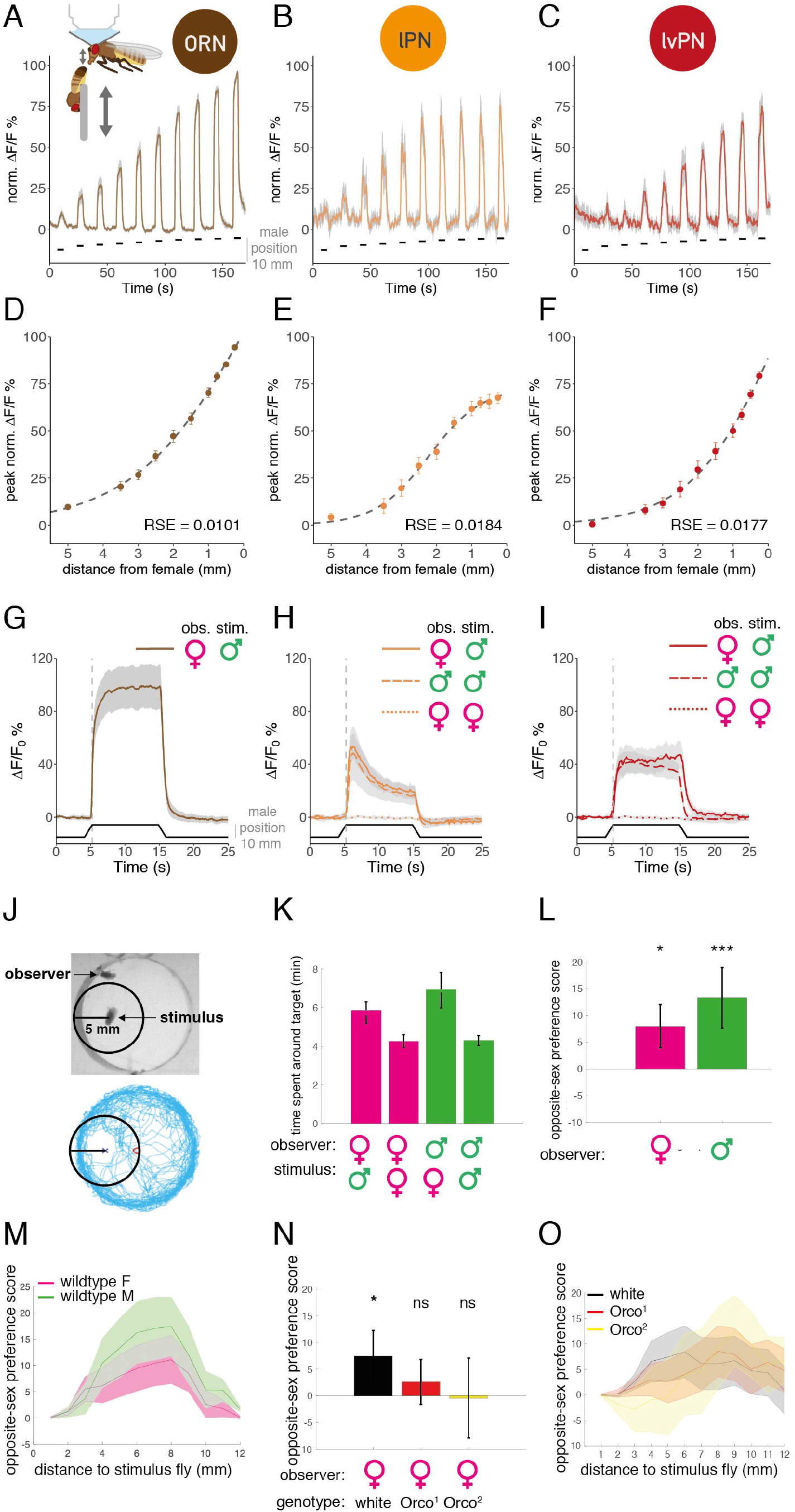
Parallel cVA pathways differentially signal male distance and sustained presence. **A, B, C**: GCaMP6f responses to a male at ten distances (see methods) in ORN (B, n = 10), lPN (C, n = 9) and lvPN (D, n = 10) axons. Shaded area is SEM. Black lines mark the time of male presentation, right y axis corresponds to the distance from the starting position. **A inset**: Experimental setup for *in vivo* 2-photon imaging and male presentation. The distance is measured between the male’s abdomen and the observer fly’s antennae. **D, E, F:** distance response curves in ORN (F), lPN (G) and lvPN (H) based on B, C, D respectively. y axis: peak values of normalized traces at the ten distances from all measured flies, 3 presentations, error bars are SEM. Dashed line shows the best sigmoidal fit: residual standard error (RSE) and half maximal distance (ED50) were: (F) RSE = 0.0101, ED50 = 2.2 mm, (G) RSE = 0.0184, ED50 = 2.4 mm, (H) RSE = 0.0177, ED50 = 1.5 mm. **G:** GCaMP6f responses in ORN axons to 10 s male presentation (0.75 mm), female fly imaged. Average response from 10 flies, 6 presentations, gray area is SEM. Bottom black trace shows male position. **H, I:** GCaMP6f responses in females to a male stimulus (solid line), in males to a male stimulus (dashed line) and in females to a virgin female stimulus (dotted line) in lPN (H) and lvPN (I) axons. Presentation same as in G. Quantification in Figure S2C. **J**: Top: A representative video frame with a stationary fly (“stimulus”), and a freely behaving fly (“observer”). The black circle shows 5 mm around the male. Bottom: trajectory of the observer over 20 minutes. Body centers were tracked. **K:** Time spent by an observer female (magenta) or male (green) within 5 mm from a stimulus fly (female or male), during 20 min recordings. Females with a female stimulus: 4.25 min; with a male stimulus: 5.84 min. Males with a male stimulus: 4.29 min; with a female stimulus: 6.95 min. Legend continued on next page. **L:** opposite-sex preference score (OSP): wildtype male and female flies spend more time within 5 mm radius of a stimulus fly from the opposite sex. 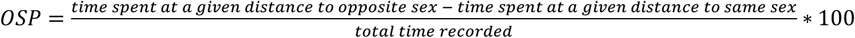 Female OSP=7.95±4, male OSP=13.29±5.6. **M:** OSP at increasing distances from stimulus: for both wildtype males and females, OSP increased with increasing distance from stimulus until 8 mm then started decreasing. Lines represent mean OSP within cumulative 1 mm bins. Shaded area is SEM. **N:** *Orco^1^* and *Orco^2^* females have impaired OSP within 5 mm from a male stimulus. *OSP(white)*= 7.44±4.8; OSP(*Orco*^1^) = 2.61 ±4.1; OSP(*Orco*^2^) = 0.72±7.4. **O:** OSP at increasing distances from stimulus: *Orco^1^* and *Orco^2^* females shifted their OSP to greater distances from a male stimulus. Lines represent mean OSP within cumulative 1 mm bins. Shaded area is SEM. Throughout the figure, ns p>0.05, * p<0.05; *** p<0.001. See genotypes and statistics in Supplementary File 1.

We tested ten positions, returning to a starting point 10 mm away between each step. All three cell types showed highly reliable activity levels based on male position, indicating that cVA concentration at the antenna can signal male distance (Figure 2A, B, C). We described the distance tuning with a sigmoidal function of the maximal response at each position (Figure 2D, E, F). ORNs already respond reliably at 5 mm distance with the widest dynamic range; responses continue to increase as distances decrease. In contrast, the lPN tuning curve is sigmoidal, plateauing at 1 mm. lvPNs respond more weakly to mid-range distances (5-2 mm) but their response grows sharply and without saturation from 2-0.25 mm. lPNs are therefore more cVA-sensitive than lvPNs, consistent with their greater number of ORN synaptic inputs (Figure 1A, Figure S1F).

We next assessed PN adaptation by keeping the stimulus male at 0.75 mm from the imaged fly’s antennae for 10 s. ORN responses reach their maximum more slowly than lPNs, and lPNs adapt more strongly during the stimulus (Figure 2G, H) consistent with results for other glomeruli (Bhandawat et al., 2007). Interestingly, DA1 lvPNs showed no adaptation in axonal calcium signal throughout the 10 s stimulus (Figure 2I). To confirm the cVA-specificity of these responses, we repeated these experiments with a virgin female stimulus; 10 s presentations elicited no responses in either lPNs or lvPNs (Figure 2H, I, Figure S2B), demonstrating that neither movement artifacts nor virgin female odors activate these PNs, in line with previous results for lPNs (Clowney et al., 2015; Dweck et al., 2015). However, for the same stimulus, lPNs and lvPNs respond similarly in males and in females (Figure 2H, I, Figure S2C).

Other fly odors have been proposed to act as short range attractive pheromones (Dweck et al., 2015) but their spatial receptive fields have never been measured directly (Agrawal et al., 2014; Root et al., 2008). Our results suggest that cVA on a male fly can only be detected by another fly when they are less than 5 mm (two body-lengths) apart. To test the behavioral significance of this detection range we monitored (under long wavelength infrared illumination) the time that an observer fly spent within 5 mm of a decapitated, stationary stimulus fly (Figure 2J, K); we found that both virgin males (63% extra) and females (37%) spent more time near a fly of the opposite sex. For statistical analysis, we calculated an opposite-sex preference (OSP) score inside circles of increasing radii centered on the stimulus fly (Figure 2L). There was a strong preference by 5 mm which then declined at large radii indicating a loss of preference when far from the stimulus (Figure 2M, Figure S2E). We then asked whether OSP depends on olfaction using loss-of-function mutants in *Orco* (Odorant receptor coreceptor), rendering flies insensitive to most odors, including cVA (Larsson et al., 2004). *Orco* mutant females lost their OSP within 5 mm distance (Figure 2N, Figure S2D) shifting their preference to greater distances (Figure 2O, Figure S2F), suggesting that the spatial preference of females to males within cVA sensation range (5 mm distance) depends on odors sensed via Orco, in agreement with previous results (figure S8B of Sun et al., 2020).

### Male cVA can generate a detectable gradient across the two antennae

Based on the data so far, we hypothesized that cVA might enable flies of either sex to locate a male or mated female during social behaviors. We investigated this using video data from the same behavioral assay to analyze turning behavior within the range of cVA detection (see methods). We quantified the facing angle relative to a male stimulus: a positive change in facing angle corresponds to a turn away from the male (Figure 3A, Figure S3A). We found an approximately 3:1 bias to turn away from the stimulus male within the 2-5 mm cVA detection range but no bias at greater distances (Figure 3B, E). Although we expect males to avoid other males, both sexes showed similar results (Figure 3B, E), so these rapid turns may reflect collision avoidance in the complete darkness of the assay rather than a sexual behavior.

**Fig. 3.**
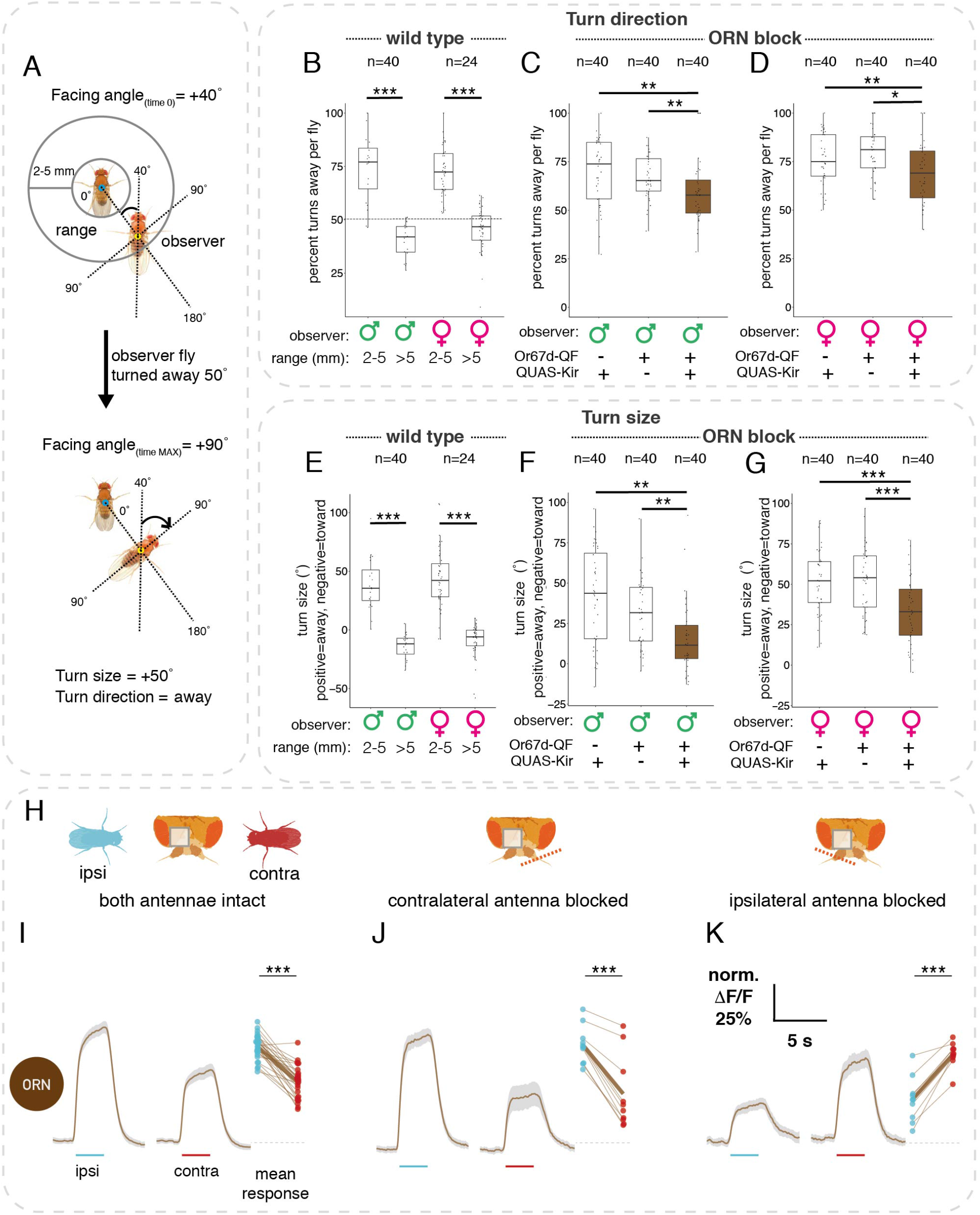
cVA sensing affects turning relative to a male and the inter-antennal distance is sufficient to detect the relevant cVA gradient. **A:** Turn size and direction calculation, using FlyTracker’s facing angle output. Turns away from stimulus male were positive, turns toward were negative. Per turn, facing angle at time 0 was subtracted from the maximal facing angle within the following 3 s. Turns inside the range are when an observer fly is within 2-5 mm away from the immobilized stimulus male. In the example shown, turn size is +50” and turn direction is away. See also Figure S3A. **B**: The percentage of turns away per wildtype fly inside or outside the range. For both males and females, turn direction inside the range is significantly different than outside. Dotted line shows 50%. **C, D:** The percentage of turns away per fly within 2-5 mm from a stimulus male is lower in Or67d-blocked males (C) or females (D). **E**: Turn size distribution in wildtype males or females inside or outside cVA range. Dots are average turn size per fly. For both females and males, turns inside the range are significantly greater than outside. **F, G**: Turn size distribution per fly while inside the range. Turns are smaller in Or67d-blocked males (F) or females (G). **H:** Antennal manipulations with respect to an imaging ROI (gray square). Left: both antennae intact, middle: antenna contralateral to ROI blocked, right: antenna ipsilateral to ROI blocked. Cyan and red flies show the position of ipsi- and contralateral male presentations, respectively. **I:** ORNs respond stronger to a male presented ipsilaterally. Left: GCaMP6f responses in ORN axons to ipsi- and contralateral male presentation. Male presentation time marked by cyan (ipsilateral) and red (contralateral) lines. Average response from 14 flies (28 hemispheres), 6 presentations, gray area is SEM. Right: mean responses of hemispheres to ipsi- and contralateral stimuli. **J:** Same as G, contralateral antenna blocked. n = 10 **K:** Same as G, ipsilateral antenna blocked. n = 10. Throughout the figure, * p<0.05; ** p<0.01; *** p<0.001. See genotypes and statistics in Supplementary File 1.

To test whether cVA is involved, we repeated the assay after blocking Or67d ORNs. For a male observer fly, the turn bias was reduced to 1.3:1 (Figure 3C) and the turn size was almost five times smaller (Figure 3F); for females, the reductions were significant but not as large (Figure 3D, G). These results indicate that cVA and DA1 circuits play a significant role in controlling rapid turns near a target male. Furthermore, our results suggest that cVA has dual actions, contributing to sex-shared instantaneous spatial orientation as well as to longer timescale sexually dimorphic behaviors.

For cVA to modulate turn direction, we hypothesized that flies compare signals from their two antennae. There is precedent for this idea in previous studies showing that flies can detect artificial odor gradients created by stimulation directed separately at each antenna (Agarwal and Isacoff, 2011; Borst and Heisenberg, 1982; Gaudry et al., 2013); however the relevance of this phenomenon to natural behavior remains unclear. We imaged Or67d ORN axons while presenting a stimulus male 1.25 mm from the observer fly (Figure S3B). We compared responses when the stimulus was the same side (ipsilateral) or opposite (contralateral) to the imaged antennal lobe (Figure 3H). ORNs were indeed more strongly activated by ipsilateral presentations (Figure 3I). Given the steep distance tuning in Figure 2D, this is what we would naively expect if ORNs projected only to the ipsilateral antennal lobe. However the situation is more complex since most ORNs in *D. melanogaster,* including Or67d ORNs, project to both sides of the brain (Stocker et al., 1983). How is the observed bilateral contrast generated given that we imaged summed responses coming from both antennae? We addressed this with antennal block experiments. When we selectively recorded ipsilateral ORNs from the same side as the imaged hemisphere (by blocking the contralateral antenna), responses to ipsilateral stimuli remained larger (Figure 3J). Similarly, when selectively imaging contralateral ORNs (by blocking the ipsilateral antenna) we saw larger responses to a contralateral male (Figure 3K). Thus bilateral contrast originates from intrinsic differences in ORN signaling levels based on stimulus distance, but we cannot rule out the possibility that they are boosted by circuit interactions at the axon terminals.

Our results show that a simple naturalistic stimulus can evoke odor responses with strong bilateral contrast based on highly sensitive detection combined with a very short range of action. Crucially, we directly demonstrate that the ~0.3 mm inter-antennal distance of flies is sufficient to detect the gradient in cVA activity from a nearby male on one side of the fly. It is noteworthy that pheromone sensitive ORNs are located on the lateral distal portion of the antenna, maximizing this distance (van der Goes van Naters and Carlson, 2007).

### Glomerulus-specific inhibition increases bilateral contrast for cVA

We have shown that a nearby stimulus male can produce bilateral contrast in sensory neuron activity. How is this bilateral ORN input processed by projection neurons (PNs)? We repeated our imaging experiments for lPNs and lvPNs in a female observer (Figure 4A), finding that ipsilateral male stimuli evoked stronger responses in both PN types (Figure 4B, E) and confirming this observation in male lPNs (Figure S4B). However, the difference between ipsi- and contralateral responses was consistently larger in PNs than ORNs (Figure 4F). Both DA1 lPNs and lvPNs receive more synapses from ipsilateral ORN axons (5184 vs 3580 for lPNs; 157 vs 113 for lvPNs). This selective pooling of ipsilateral inputs – which is typical of most PNs (Schlegel et al., 2021; Tobin et al., 2017) – provides a partial explanation for the increased bilateral contrast in PNs. However we suspected additional mechanisms were at play and therefore carried out a series of antennal block experiments to test the significance of active bilateral signals.

**Fig. 4.**
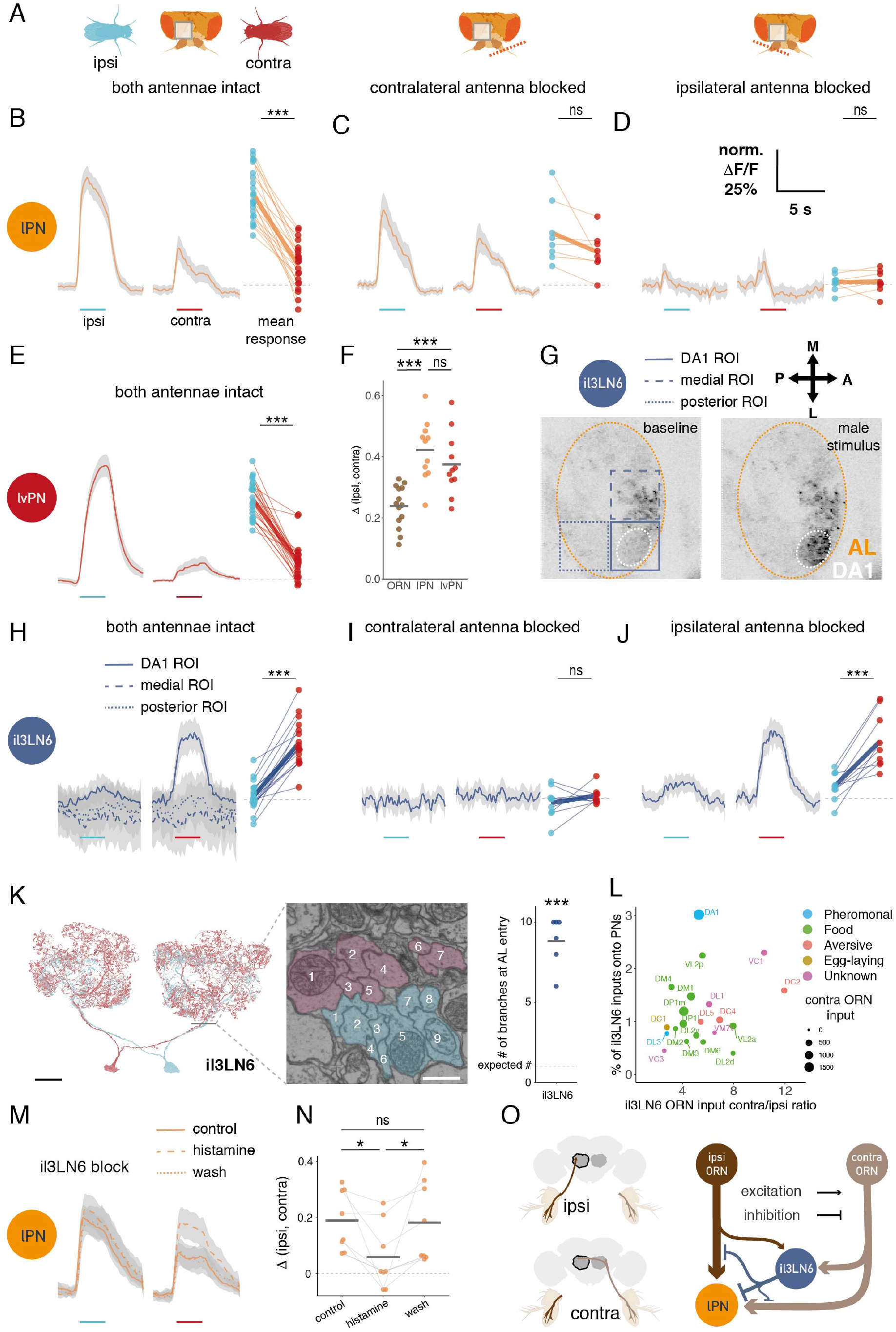
An active mechanism increases bilateral contrast in cVA sensing. **A**: Antennal manipulations with respect to an imaging ROI (gray square). Same as Figure 3H. **B**: lPNs respond stronger to a male presented ipsilaterally. Left: GCaMP6f responses in lPN axons to ipsilateral and contralateral male presentation. Male presentation time marked by cyan (ipsilateral) and red (contralateral) lines. Average responses from 11 flies (22 hemispheres), 6 presentations, gray area is SEM. Right: mean responses of hemispheres to ipsi- and contralateral stimuli.**C**: Same as B, contralateral antenna blocked. n = 8 **D**: Same as B, ipsilateral antenna blocked. n = 8 **E**: lvPNs respond stronger to a male presented ipsilaterally, stimulus as in B, intact antennae. n = 11 (22 hemispheres) **F:** Bilateral contrast is larger in both PN types than in ORNs. Bilateral contrast was calculated as the difference of mean responses to ipsi- and contralateral male presentation. **G:** Example images of il3LN6 GAL4 (VT046100) GCaMP before and during contralateral male presentation (dorsal AL). Three ROIs marked that were used to quantify the responses in different parts of the il3LN6 arbor (traces in H). Pixel gray level shows GCaMP signal intensity. **H:** il3LN6 responds stronger to a male presented contralaterally in the DA1 glomerulus, and shows no responses in adjacent arbors. Left: same stimulus as in B, intact antennae. n = 9 (18 hemispheres). Right: mean responses of hemispheres to ipsi- and contralateral stimuli. **I:** Same as H, only DA1 ROI shown, contralateral antenna is blocked. n = 9 **J:** Same as H, only DA1 ROI shown, ipsilateral antenna is blocked. n = 9 **K:** Left: EM morphology of il3LN6 neurons in FAFB, partial reconstruction. Scale bar: 20 μm. Middle: axon cross sections of two il3LN6 neurons before entering the AL. Scale bar: 750 nm. Right: number of branches entering the AL for il3LN6 neurons from the hemibrain (2) and FAFB (4) datasets. Horizontal bar shows the mean (8.83 + 1.46), which is significantly different from 1, the expected value for fly neurons. **L:** Connectivity of il3LN6 by glomerulus in the hemibrain dataset. X axis shows the ratio of contralateral and ipsilateral ORN input to il3LN6, Y axis shows the fraction of il3LN6 inputs to uniglomerular PNs, the size of the points is proportional to the number of contralateral ORN inputs to il3LN6 in a glomerulus. Five glomeruli with known bilateral ORN innervation (VA1d, VA1v, DC3, VL1, VP1d) were excluded from this analysis due to missing side information for ORNs in the hemibrain. **M:** Blocking il3LN6 decreases bilateral contrast in lPNs. lPN axon responses, same stimulus as in B, before (control), during (histamine), and after (wash) chemogenetic block of il3LN6 neurons. **N:** Quantification of M. The difference between lPN responses to ipsilateral and contralateral male presentation is smaller during il3LN6 block than before adding and after washing out histamine. **O:** Connectivity of DA1 ORNs, lPNs, and il3LN6 in one hemisphere, data from hemibrain. The line width is proportional to the base 2 logarithm of the number of synapses for a given connection. Number of synaptic connections: ipsiORN - lPN: 5184; contraORN - lPN: 3580; ipsiORN - il3LN6: 359; ipsiORN - il3LN6: 1901; il3LN6 - lPN: 465; il3LN6 - ipsiORN: 297; il3LN6 - contraORN: 165 Throughout the figure, * p<0.05; ** p<0.01; *** p<0.001. See genotypes and statistics in Supplementary File 1.

We anticipated that antennal block would decrease the amount of sensory input and therefore PN responses, but that this reduction might differentially affect signals from the two antennae. Instead we found that blocking an antenna actually increases PN responses in some stimulus configurations, directly indicating the presence of contralateral inhibition. For example when presenting the stimulus male on each side of the observer, blocking the contralateral antenna decreased responses to ipsilateral presentations and increased responses to contralateral presentations (compare Figure 4C with Figure 4B, Figure S4E); both effects combined to decrease the bilateral contrast in lPNs from 42% ΔF/F_0_ difference to 11%. Blocking the ipsilateral antenna decreased and shortened the activation compared to control, and contralateral excitation was followed by sustained decrease in lPN activity (Figure 4D). Next we presented the male centrally (as in Figure 2G) while blocking the antenna to activate only one side. In this case 1PN responses were reduced by blocking the contralateral antenna, while blocking the ipsilateral antenna caused an even more pronounced decrease below baseline than we saw with presentations on the fly’s left or right (Figure S4C, D). These data can be explained by a contralateral inhibition mechanism: when both antennae are intact, contralateral input provides both excitation (via ORNs) and inhibition onto lPNs. For an ipsilateral stimulus, the net effect is excitation, so that the response is smaller when the contralateral antenna is blocked. For a contralateral stimulus the net effect on lPNs is inhibition: blocking the contralateral antenna releases this inhibition so the response is larger. Blocking the ipsilateral antenna is in line with this model: the ipsilateral stimulus evokes a weak excitation, while the contralateral stimulus evokes a brief excitation followed by tonic inhibition.

We identified a likely source of contralateral inhibition through connectomic analysis of the fly olfactory system (Schlegel et al., 2021). il3LN6 is a large pair of local neurons that innervate both antennal lobes and arborize in ~30 glomeruli, including DA1 (Figure 4K, L). This GABAergic (Figure S4G) inhibitory neuron makes synapses onto PNs and importantly receives strongly biased ORN input: contralateral ORNs provide 5 times more synapses than ipsilateral ones (Figure 4O, L). When we inspected the EM morphology of il3LN6 neurons we found that very unusually il3LN6 splits into about 9 co-fasciculated branches before entering the AL (Figure 4K). This results in multiple smaller diameter axons increasing electrotonic separation of glomerular sub-arbors and suggests that il3LN6 is highly compartmentalized. Indeed, when we imaged il3LN6 responses to a male fly, activation was specific to the DA1 glomerulus; adjacent parts of the arbor in the imaging plane did not respond (Figure 4G, H).

il3LN6 responses to male stimuli on each side of the head showed opposite behavior to ORNs and lPNs: contralateral stimulation evoked larger responses (Figure 4H); this is consistent with strong contralateral ORN input (Figure 4L, O). Blocking the contralateral antenna completely abolished il3LN6 responses (Figure 4I), indicating that only contralateral ORN input is functional in our stimulation protocol. In line with this, blocking the ipsilateral antenna had no effect on il3LN6 (Figure 4J). These results suggest that il3LN6 inhibits lPNs when presented with a contralateral stimulus, thereby increasing bilateral contrast in lPN responses, completely consistent with the effect of antennal block on lPNs (Figure 4B, C, D, Figure S4E, F). To further test this idea, we chemogenetically blocked il3LN6 neurons by misexpressing the histamine-gated chloride channel Ort (Liu and Wilson, 2013) while measuring lPN responses to bilateral male presentation. Blocking il3LN6 with histamine reduced the difference in lPN responses between ipsi- and contralateral male presentation (Figure 4M, N), demonstrating that il3LN6 significantly increases bilateral contrast in DA1 lPNs.

DA1 lvPNs also show large differences to ipsi- and contralateral male presentation (Figure 4E), although they receive only 3 synapses from il3LN6 across the 3 lvPNs. Based on the connectivity, it is unlikely that inhibition from these LNs would directly affect lvPN responses. However, lPNs form dendro-dendritic connections with lvPNs, potentially contributing to the observed large bilateral response differences.

il3LN6 has extensive arbors so its effect on the pheromone glomerulus DA1 is unlikely to be unique. However, earlier results for another glomerulus (DM6) ruled out a contribution of GABAergic inhibition to the preference for ipsilateral ORN stimulation (Gaudry et al., 2013). To assess the broader impact of il3LN6 across all olfactory glomeruli, we compared the ratio of contra- and ipsilateral ORN inputs to il3LN6 and the fraction of inputs from il3LN6 onto canonical uniglomerular PNs (Figure 4L). Both measures should scale with il3LN6’s efficacy to provide contralateral inhibition. DA1 lPNs receive the highest proportion of their inputs from il3LN6; DM6 is weakly innervated but in e.g. DC2 and VC1 the ORN contra-ipsi bias is stronger than in DA1. This suggests that il3LN6 could have a similar role in other glomeruli, enhancing bilateral contrast of selected odors. We hypothesize that this might be of particular significance for short range, low volatility odorants.

### Projection Neurons encode male angular position

Our previous experiments maximized cVA concentration differences between the two antennae by presenting stimuli at 90 degrees to the left or right of the imaged fly. The strong bilateral contrast in those experiments, the existence of specialized local circuitry and the effects of cVA on the direction of rapid turn behavior all suggest that flies might be able to decode the angular position of another fly. We therefore presented a stimulus male at sixteen positions defined by a hexagonal lattice sampling a 2 mm radius around the imaged fly (Figure 5A, methods). We simultaneously imaged lPN dendrites on both sides of the brain, and found that responses show a spatial gradient with strongest responses when the male is nearest (1 mm) and slightly ipsilateral with respect to the imaged PN (Figure 5A). We then calculated mean responses for eleven angles at 1.7 mm distance (see methods). Left and right lPNs show symmetric angular tuning: responses were larger for stimulation ipsilateral to the imaged PN, and identical for both sides when the male was in front of the fly (Figure 5B).

**Fig. 5.**
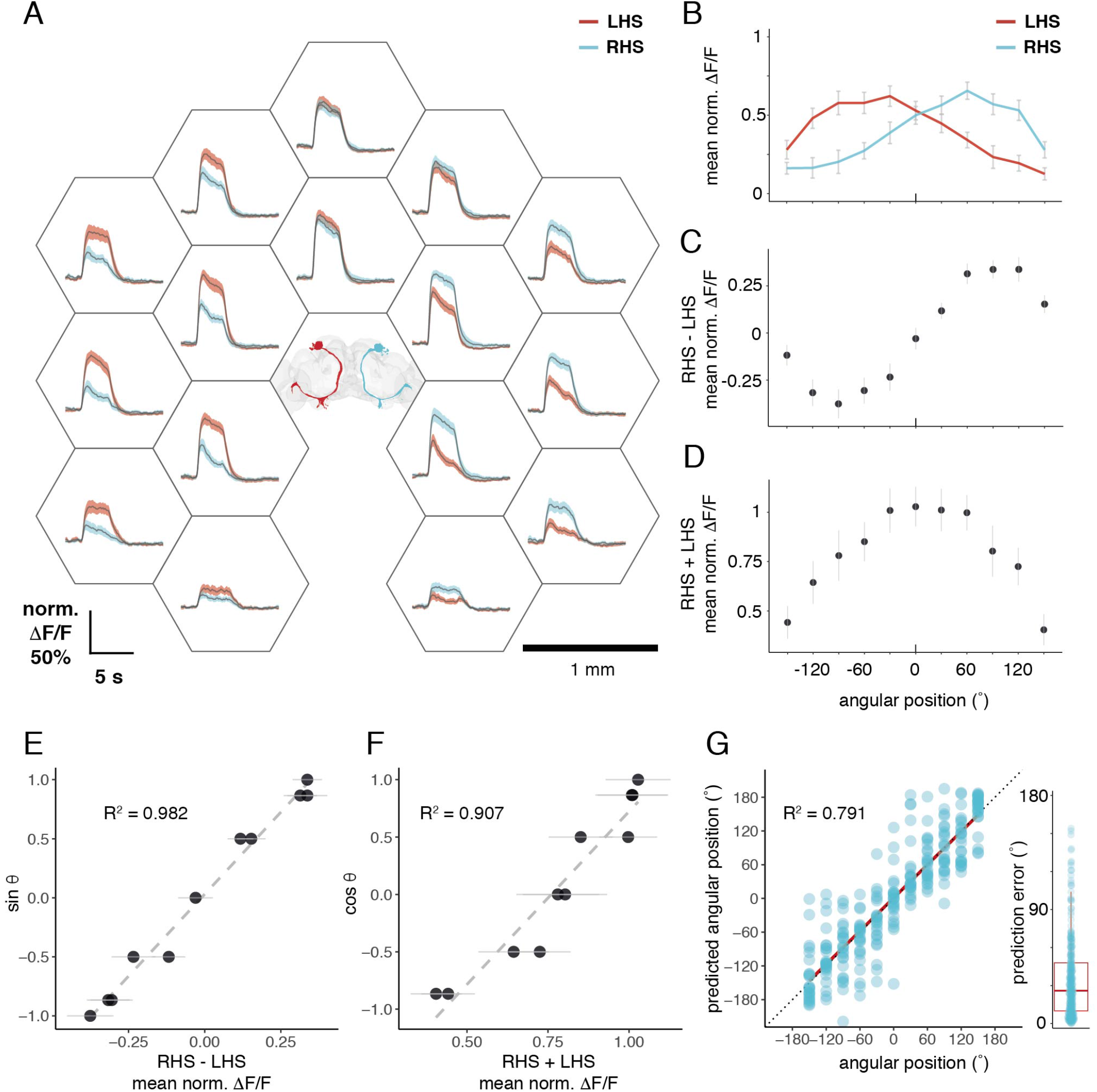
Projection Neurons encode male angular position. **A**: GCaMP6f responses of left hemisphere (LHS, red) and right hemisphere (RHS, cyan) lPN dendrites to male presentations. Positions of the male during the stimulation protocol are indicated by the center points of the hexagons around the respective responses. Positions are at 1, 1.732, and 2 mm distances, and at 30° steps in the range between −150° to +150° angular positions; the fly faces 0°, negative angles are on the left, positive angles on the right side. The position of the imaged female fly is indicated by the brain volume in the center (top view), lPNs colored based on their side as the corresponding GCaMP traces. Average response from 8 flies, 3 trials, shaded area is SEM. Scale bar: 1 mm. **B:** Angular tuning curves of left (red) and right (cyan) lPNs based on A. Six of these positions are direct measurements, five of these (at angles −120°, −60°, 0°, +60°, +120°) are based on linear interpolation from responses at 1 and 2 mm at these angular directions (see Methods). Error bars are SEM. **C:** The difference of mean right and left lPN responses at given male angular positions. Error bars are SEM. **D:** The sum of mean right and left lPN responses at given male angular positions. Error bars are SEM. **E:** The difference of right and left lPN responses correlates with the sine of the male angular position. R^2^ = 0.982, dashed line shows linear fit. **F:** The sum of right and left lPN responses correlates with the cosine of the male angular position. R^2^ = 0.907, dashed line shows linear fit. **G:** A linear model predicts male angular position based on bilateral lPN responses. Model formula in Wilkinson notation: (x, y) ~ (IPN_R_ - IPN_L_)+(IPN_R_ + IPN_L_). Where x and y are the Cartesian coordinates of the male fly, and IPN_R_, IPN_L_ are right and left lPN responses, respectively. The predicted angular position is calculated from the x and y predictions for single trials from (A). Predicted male angular positions correlate with the actual angular position, R^2^ = 0.792. Red solid line shows the linear fit, the dotted line shows x = y. Right: distribution of prediction error in angles, mean = 35°, median = 26°.

Could the fly infer a male’s angular direction by using lPN responses from both sides of the brain? We found that the difference and the sum of the left and right responses strongly correlate with the sine and the cosine of the male’s angular position (Figure 5C, D, E, F). The sine and the cosine together give a unique solution to angular position around a full rotation of a circle. The difference and the sum of bilateral lPN responses therefore provide measures to predict a male’s angular direction. Moreover, the sine and cosine of an angle are equal to the Cartesian x and y coordinates of a point on the unit circle. We devised a bivariate linear model using lPN responses from the two hemispheres, where the difference and the sum of the right and left lPN responses are used as predictors, and the male’s x and y position are the response variables. This model accurately predicts stimulus position when given the response of left and right PNs (Figure S5A, B, median error 1.3 mm). For angular direction the median prediction error was 26° (Figure 5G).

We found an inverse relation between signal intensity and distance that is typical of most sensory systems (see also Figure 2). However intensity, the amount of cVA on males, depends on mating history and age (Kuo et al., 2012). While this creates ambiguity in estimating male distance, our model shows that angular direction can be computed by comparing lPN responses from both sides of the brain which is reliable across different cVA concentrations. Angular direction is most important for orientation behavior. Furthermore, flies likely use temporal correlations in the olfactory signal and other sensory modalities (e.g. vision) to improve distance coding.

### cVA PNs target a large and diverse array of third-order neurons

DA1 lPNs and lvPNs are the only two uniglomerular, excitatory PNs relaying cVA signals from the antennal lobe to higher brain centers. How is cVA information then read out by downstream circuits? We carried out comprehensive connectomic analysis of third-order targets; in contrast to the limited divergence at the first synapse, we found a large and diverse set of downstream targets (Figure 6A, B, Supplementary Video S1). We classified these neurons based on their input selectivity (DA1 selective, mixed olfactory, multimodal) and their projection patterns (LN, ON, DN: local, output–i.e. connecting different neuropils–or descending neurons projecting to the nerve cord); we (Dolan et al., 2019) or predictions using synaptic ultrastructure information (Eckstein et al., 2020). lPNs synapse onto 40 downstream cell types, lvPNs onto 11 (see methods for inclusion criteria, Figure 6A); only 4 cell types are shared. While the majority of lPN target output neurons receive a mix of olfactory inputs (18) or integrate odors and other sensory channels (10), there are 3 cholinergic output neurons that are DA1 selective. lvPN targets were mostly multimodal (7) with only a few olfactory downstream partners (2) consistent with lvPN projections to the multimodal SIP area. This diversity of third-order cell types, ranging from DA1 selective to multimodal integrators could allow the representation of distinct features of a single stimulus.

**Fig. 6.**
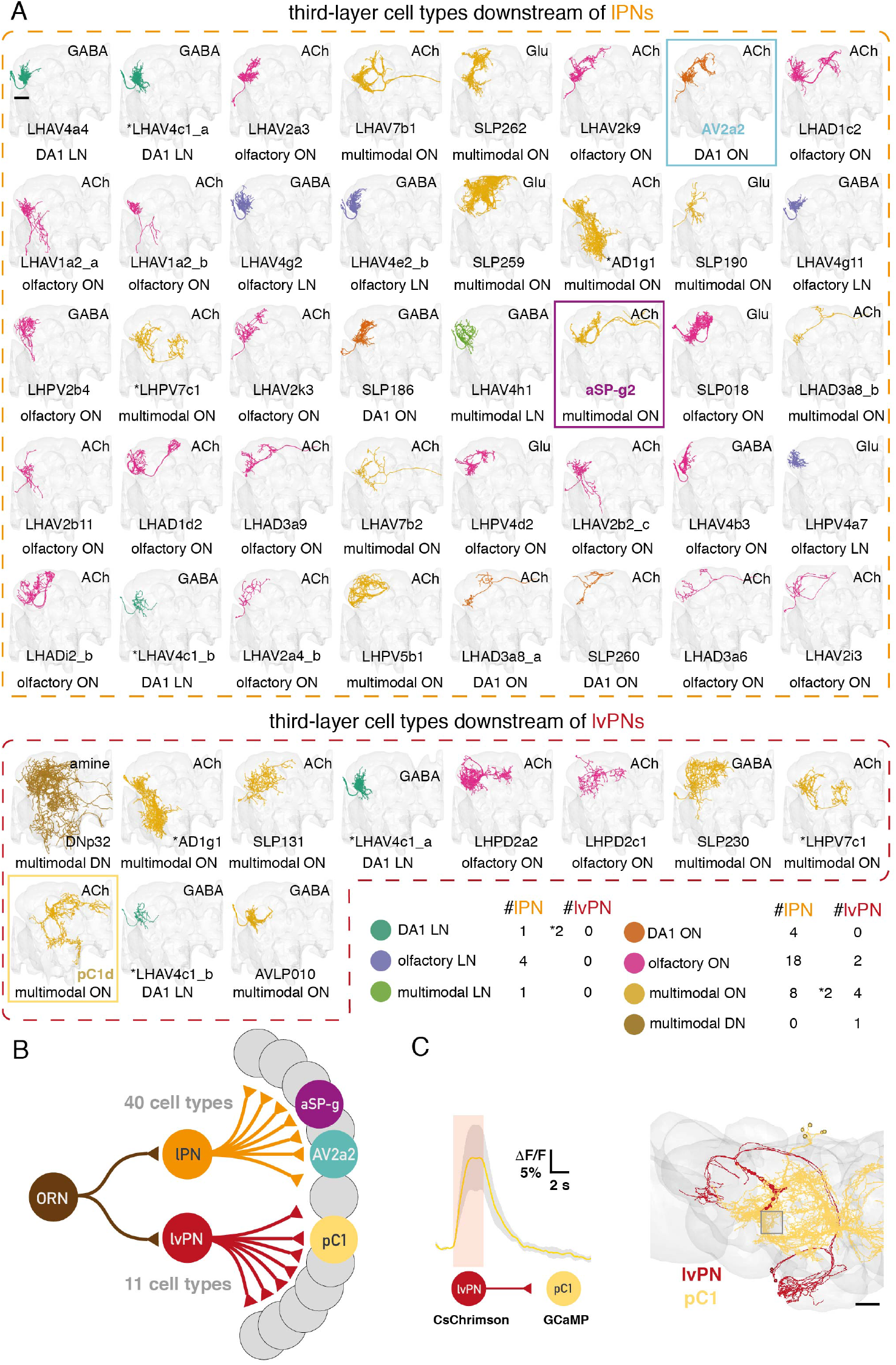
cVA PNs target a large and diverse array of third-order neurons. **A:** Downstream cell types of DA1 lPN (top) and DA1 lvPN (bottom) in the hemibrain, ordered by the absolute number of inputs from the respective PN type. See methods for inclusion criteria. Number of cell types per class shown in the bottom right table. Colors represent broad cell type classes. Magenta: multimodal local neuron (LN), purple: olfactory output neuron (ON), green: multimodal ON, turquoise: DA1 selective ON, orange: olfactory LN; yellow: multimodal descending neuron (DN). ACh: acetylcholine, GABA: γ-aminobutyric acid, Glu: glutamate. Scale bar: 40 μm. **B:** Circuit diagram of the first three layers of cVA processing based on A and B. **C:** Left: pC1 axons respond to lvPN optogenetic activation. Red area shows the time of optogenetic stimulation; Average response from 6 flies, 6 stimulations, shaded area is SEM. Middle: direction of the lvPN - pC1 connection. Right: EM reconstruction of lvPNs (red) and the five pC1 cells (a-e) (yellow, from Wang et al. 2020) neurons in FAFB, top view. Red circles show the location of lvPN to pC1 synapses. Gray square shows the ROI used for recording pC1 responses. Scale bar: 20 μm.

To further understand the logic of information processing we selected a subset of these third-order cell types. Downstream of lPNs we selected two cholinergic ONs: first the *fruitless+,* sexually dimorphic aSP-g neurons (Cachero et al., 2010; also called aSP8, Yu et al., 2010), that were also found in earlier work from our lab (Kohl et al., 2013) and are multimodal according to our connectomic analysis; second, a previously undescribed cell type, AV2a2, as it is distinct from aSP-g. AV2a2 are sexually isomorphic and importantly, DA1 selective–the labeled line nature of the stimulus is kept in the third layer. Downstream of lvPNs we chose the *doublesex+* sexually dimorphic pC1 neurons, as their effect on female sexual receptivity is similar to what we observed with lvPNs (Figure 1F, G). The direct lvPN - pC1 connection suggests that this shallow circuit is responsible for the cVA responses observed in pC1 neurons. Given that lvPN inputs only make up a small fraction of all pC1 inputs (1.0%) we wanted to test whether lvPNs are sufficient to activate pC1 neurons. Indeed, optogenetic activation of lvPNs generated calcium responses in pC1 neurons (Figure 6C). Anatomy, neural responses, and behavioral data together indicate that lvPNs relay cVA information to pC1 neurons, and suggest that pC1 conveys the receptivity promoting effect of lvPNs in females.

### Third-order neurons extract distinct features of a male from cVA stimuli

First, we focused on AV2a2 and pC1 (Figure 7A, B; AV2a2 driver lines in Figure S6A, B). A single presentation of a male for 10 s evoked drastically different responses in these two cell types. pC1 show a sustained and slowly decaying response, while AV2a2 selectively respond to the onset of the male stimulus (Figure 7C). Zhou et al., (2014) showed that cVA is integrated with courtship song by pC1 neurons, therefore pC1 activity might reflect a courting male’s reproductive quality. We corroborate this idea by showing that pC1 neurons respond tonically to a male’s presence, similarly to lvPNs (Figure 7C).

**Fig. 7.**
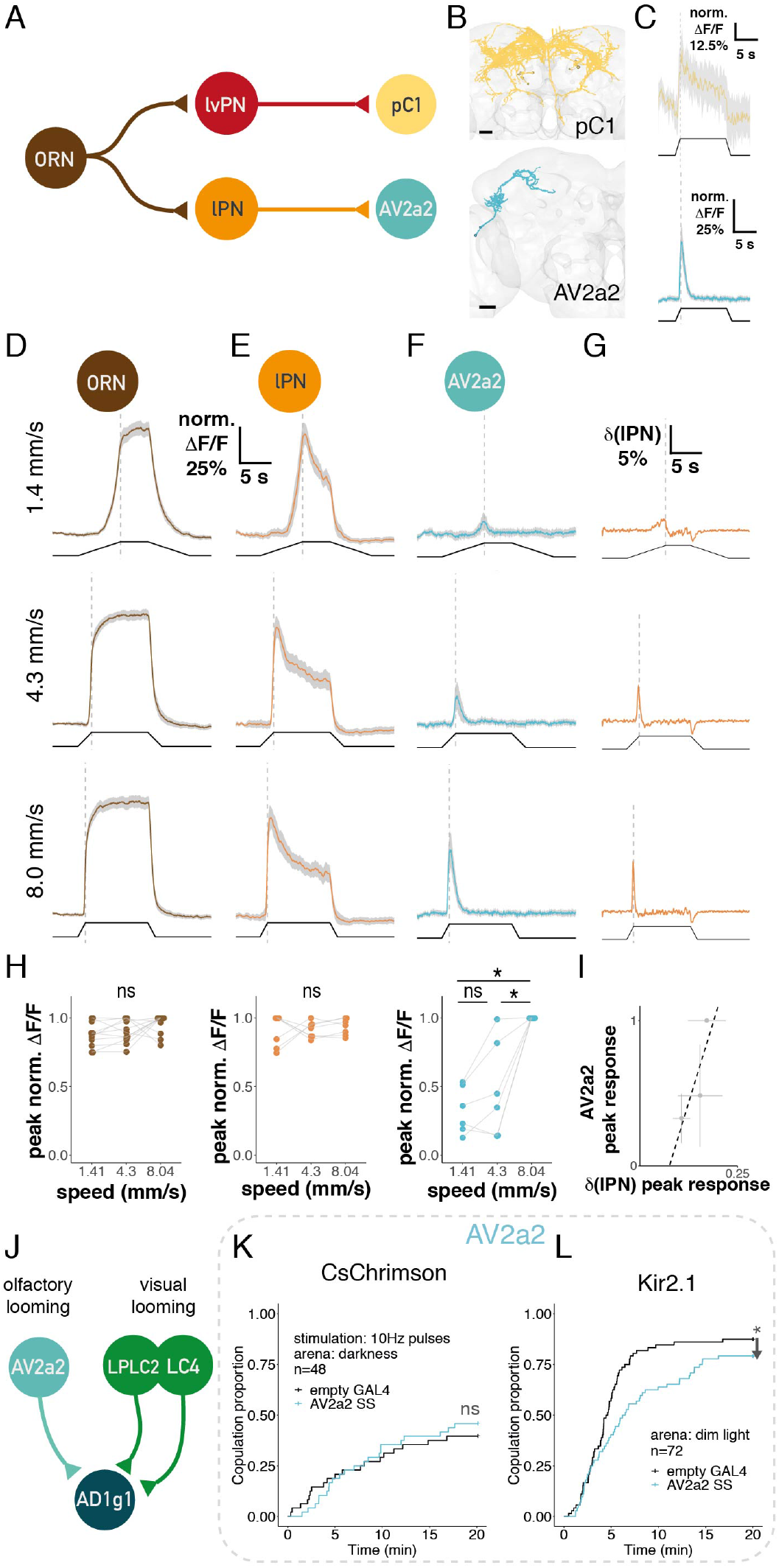
Third-order neurons extract distinct features of a male from cVA stimuli. **A:** Connectivity of DA1 ORNs, lvPNs, pC1, lPNs, AV2a2 based on hemibrain. Number of synaptic connections: ORN - lvPN: 286; lvPN - pC1: 45; ORN - lPN: 9187; lPN - AV2a2: 115 **B:** EM reconstructions in FAFB of pC1 (top, from Wang et al. 2020) and AV2a2 neurons (bottom). Scale bars: 20 μm. **C:** GCaMP6f responses in pC1 (top) and AV2a2 axons (bottom) to male presentation, same stimulus as Figure 2G. Average responses from 6 flies, 6 presentations, shaded area is SEM. **D:** GCaMP6f responses in ORN axons to male presentation (0.75 mm) at different speeds (shown on the left). Average responses from 11 flies, 6 presentations, shaded area is SEM. Bottom black trace shows male position, dashed line marks the end of the approach. **E:** GCaMP6f responses in lPN axons, stimulus as in D, n = 7. Data with the highest speed was included in Figure 2H. **F:** GCaMP6f responses in AV2a2 axons, stimulus as in D, n = 6. Data with the highest speed also shown in (C). **G:** The differential of the lPN response trace to presenting a male at different speeds, based on data in (E). **H:** ORN and lPN maximal responses are not dependent on male speed, while AV2a2 responses are; based on data in (D, E, F). Data points from individual flies connected with gray lines. **I:** lPN response differential peaks correlate with AV2a2 peak responses. Error bars are standard deviation. Dashed line shows the linear fit, R^2^ = 0.71, p = 0.36. **J:** Circuit diagram of AD1g1 input cell types. Number of synaptic connections: AV2a2 - AD1g1: 142; LPLC2 - AD1g1: 474; LC4 - AD1g1: 434. **K, L:** Manipulating AV2a2 SS in females paired with wildtype males. Curves represent the proportion of mated females over time. (Q) Pulsed optogenetic activation had no effect on female receptivity. (R) blocking with Kir2.1 decreased female receptivity. Throughout the figure, * p<0.05; ** p<0.01; *** p<0.001. See genotypes and statistics in Supplementary File 1.

The observation that AV2a2 neurons have a phasic ON-response, unlike lPNs, suggests that they might be activated specifically by fast increases in lPN activity. To test this, we varied the approach speed of the stimulus male, thereby altering the speed of cVA concentration change at the antennae while measuring responses in ORNs, lPNs, or AV2a2. In ORNs and lPNs we observed a speed dependent rise-time in intracellular calcium, but no difference in maximal responses with different approach speeds (Figure 7D, E, H). In contrast, in AV2a2 both rise-time and peak response depended on male speed (Figure 7F, H).

How is lPN activity transformed to create the responses observed in AV2a2? We found that the positive part of the differential of the lPN response trace looks qualitatively similar to the AV2a2 response traces (Figure 7G). Furthermore the size of the peaks on the lPN differential trace are correlated with the evoked peaks in AV2a2 at different speeds (Figure 7I). This correlation suggests a mechanism whereby AV2a2 integrates lPN activity on a short time-scale coupled with an intrinsic adaptation or feedback inhibition. Such mechanisms could enable AV2a2 to respond to the speed of change in lPN activity rather than absolute activity level.

The postsynaptic neuron with the most inputs from AV2a2 is AD1g1, a single lateral horn output neuron projecting to the AVLP (Figure S6C). We found that AD1g1 receives strong visual input from cell types that are selectively tuned to the size (LPLC2) or velocity (LC4) of looming stimuli (Ache et al., 2019; Klapoetke et al., 2017), Figure 7J). We therefore speculate that AD1g1 neurons integrate visual looming with an olfactory signal from AV2a2 neurons encoding male speed, and thereby AD1g1 creates a specific representation of an approaching male by multimodal integration.

To test the behavioral role of AV2a2, we used optogenetic activation or genetic ablation in female flies during a courtship assay as in Figure 1. Optogenetic activation of AV2a2 had no effect on female receptivity, while constant silencing of AV2a2 reduced female receptivity (Figure 7K, L). We therefore propose that AV2a2 activity is not a sexually decisive signal on its own, but that its suggested role in detecting male approach may be required for normal courtship. We note that the AV2a2 driver line labels a pair of bilateral neurons innervatingthe lateral accessory lobes (Figure S6B), and we cannot exclude the contributions of this cell type to the observed phenotypes.

### Multimodal integration is key to controlling female receptivity

So far we have demonstrated how multiple olfactory percepts can be generated from a single labeled line of incoming cVA information. However, cVA may have quite different meanings in different contexts: for example it is transferred from males to females during mating (Brieger and Butterworth, 1970). Our connectomics analysis identified multiple cases of integration across sensory channels and modalities; this can potentially disambiguate cVA sources with very differentethological significance. For example, Kohl et al., (2013) previously showed that aSP-g neurons respond to food odors as well as cVA and we now find that they may integrate cVA with other modalities (Figure 6A). When presenting a male fly, aSP-g neurons in females responded with phasic ON responses similar to AV2a2 (Figure S7H). However, aSP-g responses decreased with sequential male presentations, unlike in lPNs or AV2a2 (Figure S7I). Habituation to a male stimulus makes aSP-g suitable for coding stimulus novelty rather than positional features like distance or speed and the responses are likely further shaped by other sensory inputs.

To fully explore the sensory pathways upstream of aSP-g neurons we returned to the FAFB dataset since this contains the primary taste centers in the sub-esophageal zone (SEZ). We reconstructed all eleven aSP-g neurons in the left hemisphere; NBLAST morphological clustering (Costa et al., 2016) revealed three distinct subtypes (Figure S7A). We observed the same three subtypes when clustering aSP-g neurons from light microscope images from multiple brains (Figure S7C). Of these, aSP-g2 neurons have the largest proportion of dendritic arbor in the LH across all datasets (Figure S7B), and are the only aSP-g subtype that receives DA1 innervation. Kohl et al. (2013) found that only 70% of aSP-g neurons responded to cVA, likely corresponding to the 5/11 aSP-g2 neurons in the FAFB dataset. Although aSP-g neurons do not receive lvPN input, we found inputs from another lateroventral PN type; these lvPN2 neurons are multiglomerular and besides DA1 receive most input in the DC3 and VC4 glomeruli (Figure S7E) which are known to respond to fruit odors (Münch and Galizia, 2016). This provides an anatomical explanation for the mixed odor tuning of aSP-g (Kohl et al., 2013).

We used a sampling procedure to identify non-olfactory inputs to aSP-g neurons. We found several taste PN types that synapse onto all aSP-g subtypes (Figure S7D, E). We called the neuron providing the highest proportion of inputs to aSP-g2 (4.8%) G2N-SLP1 (gustatory second-order neuron; Figure 8A, B). G2N-SLP1 in turn receives inputs from two types of gustatory receptor neurons (GRNs). One is a labellar GRN located on the mouth parts and provides 8.9% of all inputs (lGRN, Figure 8A, B); the second is an internally located pharyngeal GRN, providing 20.4% of all inputs (pGRN, Figure S7E). aSP-g2 neurons are therefore third-order in both olfactory and gustatory pathways, serving as a point of sensory convergence.

**Fig. 8.**
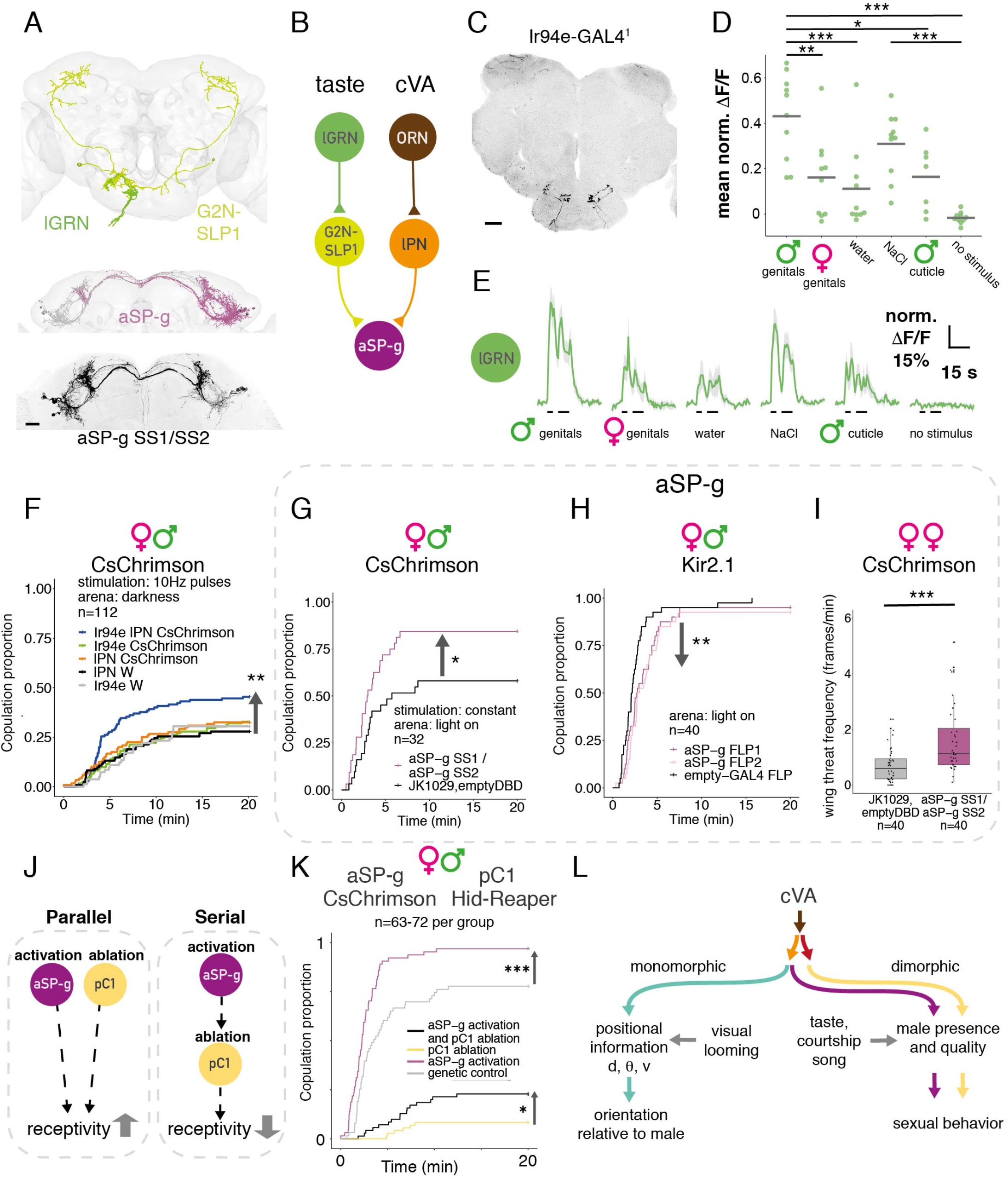
Multimodal integration is key in controlling female receptivity. **A**: Top: EM reconstruction of labellar GRNs (lGRN, green) and G2N-SLP1 (yellow) in FAFB. Middle: EM reconstruction of aSP-g neurons in FAFB, right hemisphere: purple, left hemisphere: gray. Bottom: Confocal image of aSP-g neurons in a female brain, reporter expression driven by aSP-g SS1/SS2, maximum projection. Scale bar: 20 μm. **B:** Schematic connectivity of aSP-g input pathways based on FAFB and the hemibrain. Number of synaptic connections: lGRN - G2N-SLP1: 63, G2N-SLP1 - aSP-g: 139, ORN - lPN: 8764, lPN - aSP-g: 81 **C:** Confocal image of Ir94e labellar GRNs in a female brain, reporter expression driven by Ir94e-GAL4^1^, maximum projection. Scale bar: 20 μm. **D:** Ir94e GRNs respond strongest to compounds on male genitals. Normalized GCaMP7f mean responses to labellar presentation of stimuli. Horizontal bars show the population mean of 10 flies, single presentations. **E:** Ir94e GRNs GCaMP7f responses, average responses from 10 flies. Black bars mark the time of labellar stimulation. Shaded area is SEM. **F:** Pulsed optogenetic activation of both cVA olfactory and Ir94e gustatory PNs simultaneously increases female receptivity, while individual activation does not. Activating DA1 lPN SS alone (orange), IR94e GAL4 alone (green) or both (blue) in females paired with wildtype males, compared to controls (black, gray). **G, H:** Manipulating aSP-g in virgin females paired with wildtype males. (G) Constant optogenetic activation of aSP-g SS1/SS2 increased female receptivity. (H) Using Kir2.1 to block aSP-g FLP1 or aSP-g FLP2 decreased female receptivity. **I:** Constant optogenetic activation of aSP-g SS1/SS2 in pairs of females in the light increased female-female aggression. See also Figure S1D, Supplementary Video 2. **J:** possible interactions between aSP-g and pC1 neurons: in a parallel architecture, both populations control receptivity independently, and activating aSP-g while ablating pC1 could increase receptivity compared to no activation. In a serial architecture, aSP-g effect on receptivity depends on intact pC1 and their activation cannot overcome pC1 ablation. **K:** Behavioral epistasis: optogenetic activation of aSP-g while pC1 neurons are ablated. Constant optogenetic activation of aSP-g in females either alone (purple) or while pC1 SS neurons were ablated (black). aSP-g activation increased female receptivity compared to genetic control (gray) and activating aSP-g while pC1 neurons are ablated partially rescued the receptivity impairment of pC1 ablated females (yellow). **L:** Circuit model of feature separation of an olfactory stimulus. cVA signal diverges into two parallel second-order pathways; third-order neurons represent distinct cVA-related scenarios by specific response kinetics and the integration of signals from other sensory modalities. d = distance, θ=angular direction, v = speed. Throughout the figure, * p<0.05; ** p<0.01; *** p<0.001. See genotypes and statistics in Supplementary File 1.

We focused our functional analysis on labellar inputs, since they are more likely activated by external cues. By morphological similarity we matched the labellar GRNs to Ir94e-GAL4 (see also Engert et al., 2022; Jaeger et al., 2018; Koh et al., 2014; Sánchez-Alcañiz et al., 2018). To identify candidate ligands, we imaged Ir94e GRN calcium responses to labellar stimulation in virgin female flies. As shown previously, water and NaCl activated Ir94e (Jaeger et al., 2018), but we found that presenting a male fly’s genitals evokes a larger response (Figure 8D, E). In contrast, responses to female genitals and male dorsal cuticle were similar to the smaller water responses. Compounds on male genitals, potentially contact pheromones, are therefore strong ligands for Ir94e and may contribute to female receptivity. To test this we used our courtship assay while activating Ir94e GRNs via CsChrimson in virgin females. As for DA1 lPNs, manipulating Ir94e GRNs did not change female receptivity. However, simultaneously activating Ir94e GRNs and DA1 lPNs did increase female receptivity, compared to either activation alone (Figure 8F, Figure S7J).

Ir94e gustatory and cVA pheromone signals converge on aSP-g dendrites. We directly manipulated the activity of aSP-g neurons in a courtship assay by activating or blocking neurons with multiple driver lines targeting all three aSP-g subtypes (Figure 8A, Figure 7SA, see methods). Activating aSP-g neurons increased female receptivity (Figure 8G, Figure S7K) while blocking aSP-g resulted in a small but significant decrease (Figure 8H, driver lines in Figure S7F). aSP-g therefore bidirectionally regulates female receptivity, similar to lvPNs and pC1; and activating aSP-g neurons phenocopies simultaneous stimulation of their DA1 lPN and Ir94e inputs, providing direct evidence for the behavioral significance of multimodal integration. Additional experiments showed that aSP-g does not control receptivity in mated females (data not shown) but also regulates aggression in virgin females (Figure 8I, Supplementary video S2, Figure S7L, see Figure S1D for quantification).

We can rationalize the large number of third-order cell types from a sensory coding perspective: each can be selective for a range of stimulus configurations with different ethological relevance. Like the different sensory streams converging on aSP-g, it is likely that these third-order populations interact combinatorially to control distinct behaviors. To begin testing this idea we devised a behavioral epistasis experiment in which aSP-g neurons were activated, while pC1s were genetically ablated, testing a serial vs parallel architecture (Figure 8J). As expected, ablating pC1 neurons alone suppressed female receptivity, and aSP-g activation alone increased female receptivity. In the epistasis genotype we saw a significant increase in receptivity compared to the pC1 ablated condition. aSP-g neurons can therefore partially restore female receptivity without functional pC1 neurons (Figure 8K, Figure S7M), indicating a parallel architecture (Figure 8J). This behavioral result is consistent with connectivity: aSP-g2 is not strongly connected to pC1 in the hemibrain either directly (17 synapses across five pC1), or via intermediates; furthermore they have very few common downstream partners. Connectomics analysis may therefore also be a useful guide to probing the combinatorial basis of behavior across neuronal cell types.

## Discussion

This work reveals the circuit logic by which a pheromone is used to represent qualitative and positional features separately to guide specific social behaviors. First, we show that cVA information reaches higher-order brain regions via two separate excitatory PN populations with distinct temporal dynamics. This is highly reminiscent of coding differences in mitral and tufted cells of the olfactory bulb (Fukunaga et al., 2012) however we show that these two pathways have distinct behavioral effects, something which remains unclear in mammals. DA1 lPN manipulations did not convey the previously described behavioral effects of cVA on female receptivity or male aggression, however the DA1 lvPN pathway did. Finding a direct connection from lvPNs to pC1 revealed a surprisingly shallow circuit, where a central integrator node is reached just two synapses downstream of Or67d ORNs. The direct influence of olfaction on circuits organizing behavior is also apparent in mammalian brains, where odor information reaches the piriform cortex and the amygdala without a thalamic relay.

Second, we show that the *Drosophila* olfactory system is extremely sensitive to the position of a male stimulus fly at mm ranges (Figure 2) and that cVA sensory neurons control orienting behavior (Figure 3). This may be used by females during different courtship contexts, e.g. for turning away from a male to escape (Arez et al., 2021), or to bend the female’s abdomen towards the male (Ning et al., 2022). Males likely also use cVA-based spatial information to localize other males during fighting. Fly social interactions are most common in low light at dawn and dusk; furthermore they cannot visually distinguish males and females (Agrawal et al., 2014) so in a crowded environment, visual, olfactory, and other spatial information are likely integrated to track the position of nearby flies.

We show that cVA has a very short range of action, so that bilateral comparison of PN activity can signal a male’s angular position. In our model, using both the difference and the sum of DA1 PN responses allows unambiguous decoding of the angular position of another fly (Figure 5). In contrast, wind direction sensing relies only on the difference of antennal displacement in flies (Suver et al., 2019). Bilateral comparison of auditory stimuli is also used for prey localization in the barn owl (Knudsen and Konishi, 1979). Owls synthesize this information in higher-order auditory neurons with spatial receptive fields (Pena et al., 2001); it will be exciting to see if analogous neurons exist e.g. in the fly central complex as recently identified in mouse piriform cortex (Poo et al., 2022). However these representations are not essential: fast auditory steering in crickets depends on biomechanical rather than neural integration of lateralized signals (Hedwig and Poulet, 2004). Unilateral AV2a2 could provide such steering instructions.

Third, we show that contralateral inhibition enhances bilateral contrast in the DA1 glomerulus (Figure 4). Previous work had emphasized ipsilateral biases in sensory input to another fly glomerulus PN (Gaudry et al., 2013; Tobin et al., 2017) or suggested that contralateral inhibition might be important in both flies and mice without identifying a specific neural mechanism (Dalal et al., 2020; Mohamed et al., 2019). We show that il3LN6, a GABAergic interneuron in the AL, provides such inhibition. Although it innervates ~30 glomeruli, presenting a male fly specifically activates the arbors in the DA1 glomerulus (Figure 4G, H). This suggests that its tortuous branching allows it to act in a compartmentalized fashion, reminiscent of recent studies in larval *Drosophila* (Si et al., 2021) and the adult visual system (Meier and Borst, 2019). One might think that mixing olfactory signals from both antennae would make it harder to extract positional information. However we propose that this enables a single giant interneuron to perform efficient *local* computations in each glomerulus that would otherwise require bilateral interneurons connecting each of the 50 glomeruli. Similar considerations may drive binocular convergence in the visual system.

Fourth, we show that distinct response properties and sensory integration steps in third-order neurons create more specific representations of cVA-related scenes, allowing the flexible expression of appropriate behaviors depending on the environment. cVA is not only present on males but also on mated females (Brieger and Butterworth, 1970), on the outer layer of eggs (Everaerts et al., 2018; Narasimha et al., 2019), and in male deposits (Keesey et al., 2016; Mercier et al., 2018). Therefore, incorporating other sensory modalities (like taste in aSP-g neurons, Figure 8) and responding selectively to the temporal structure of the cVA stimulus (e.g. looming sensitive AV2a2 neurons, Figure 7) is important to establish an appropriate behavioral response. aSP-g neurons habituate strongly to male odor so appear tuned to novelty of chemosensory stimuli (Figure S7H, I); they can also act as coincidence detectors: this may happen when females encounter male deposits containing both tastants and cVA.

Compared with the phasic responses in aSP-g and AV2a2, the more sustained responses in pC1 neurons can signal the presence of a potential mate. This may transform transient sensory inputs into a longer lasting internal state–as shown in the analogous male pC1/P1 circuit (Hoopfer et al., 2015; Jung et al., 2020; Kohatsu et al., 2011). Receptivity and aggression appear to be related brain states that influence the expression of specific behaviors. It is interesting that aSP-g, like pC1, controls both states, further supporting the hypothesis that these are closely related by neuronal architecture as well as behavioral expression (Anderson, 2016).

The third-order neurons encoding more qualitative features are sexually dimorphic, whereas a positional feature, speed, is encoded by a sexually isomorphic cell type (Figure 8L). This could be a general feature of sexually dimorphic behaviors: positional circuits that can also be used in non-sexual contexts are wired similarly in male and female brains, while qualitative circuits are sexually dimorphic. This separation would favor rapid evolution of circuits selective to mating and is also consistent with recent observations of a multimodal signal that gates visually driven male courtship in flies (Hindmarsh Sten et al., 2021).

Finally we propose that odor based positional information processing shows strong similarities with other sensory modalities: bilateral comparison is used to infer angular position as in the auditory system; and positional information is split from qualitative signals to be processed separately, analogous to the *what* and *where* pathways in the primate visual cortex. Thus the fly olfactory system appears to solve similar computational and behavioral challenges with a much more compact sensory processing hierarchy than in cortex. Separate processing streams do pose a long-recognized challenge, the binding problem (von der Malsburg, 1999), in how different stimulus features can be linked. The fly is now very well-placed to provide detailed mechanistic insight into this and related problems.

## Experimental procedures

### Key Resources Table

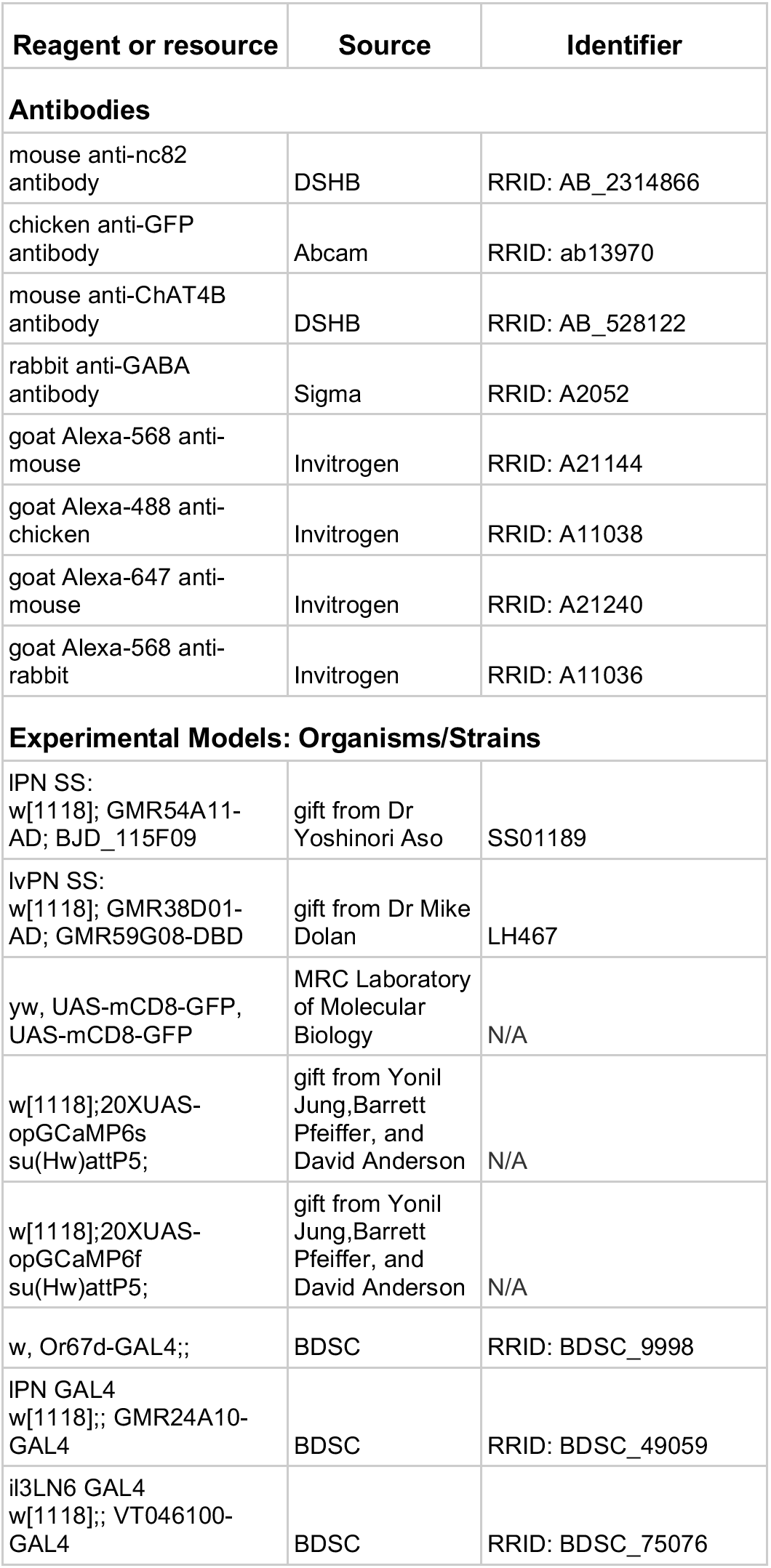
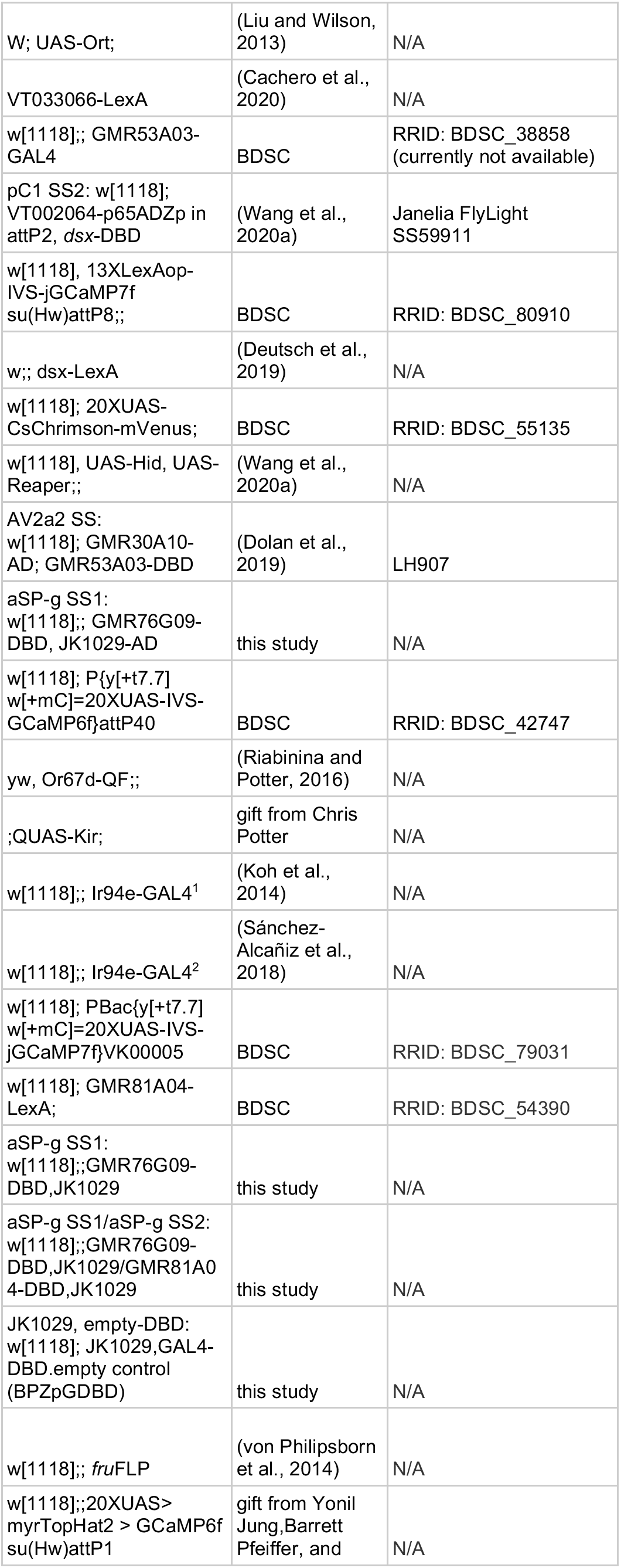
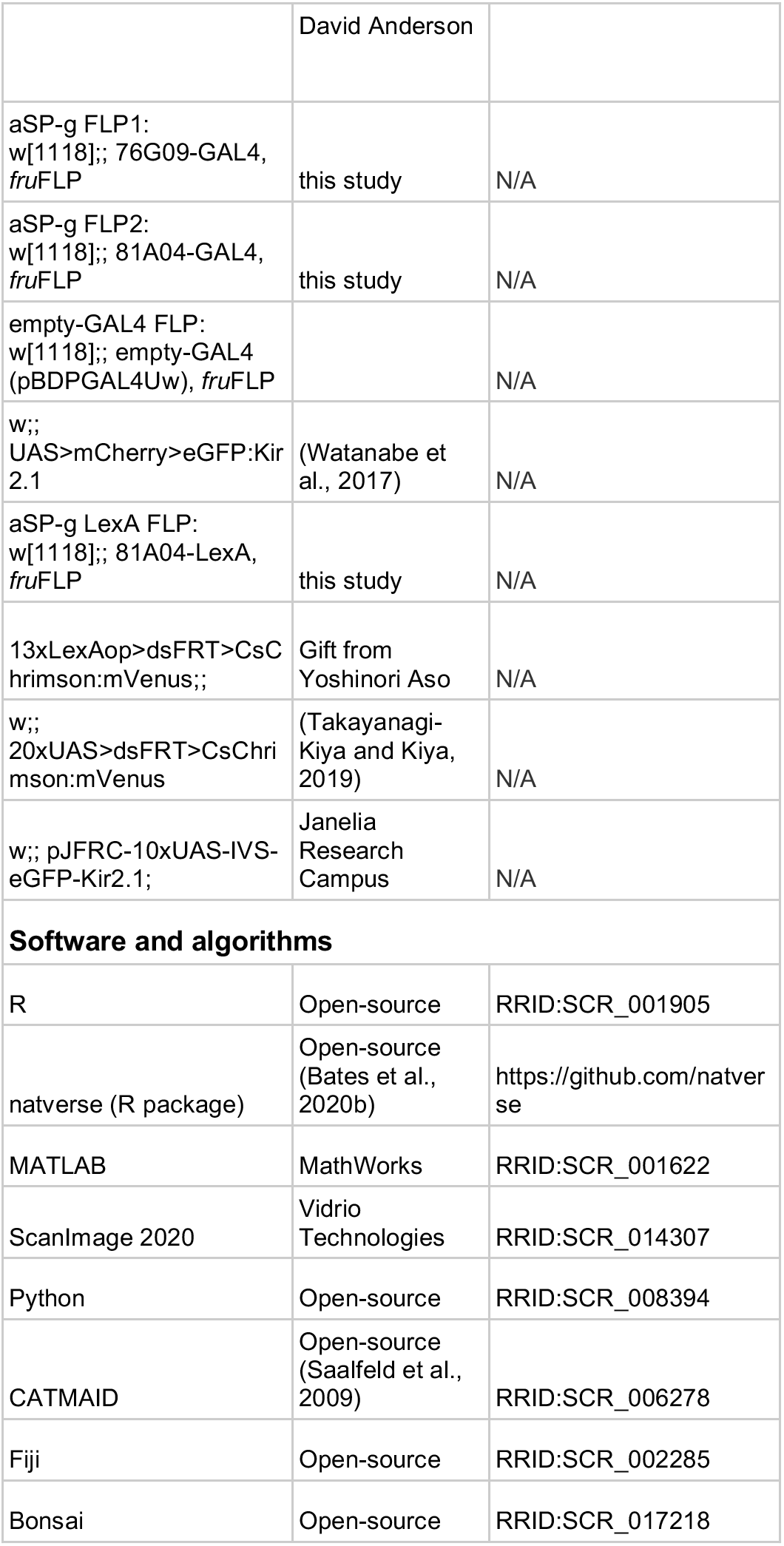

### Contact for reagent and resource sharing

Further information and requests for resources and reagents should be directed to and will be fulfilled by the Lead Contact, Gregory Jefferis).

### Drosophila husbandry

Standard techniques were used for fly stock maintenance. Flies for experiments were raised and kept at 25°C in an incubator with a 12 hour light:dark cycle, and grown on iberian Drosophila food. For optogenetic experiments the food was supplemented with 0.4 mM all-trans retinal and flies were kept in the dark.

### Split-GAL4 hemidriver combination screening

To find genetic driver lines labeling our cell types of interest our starting point was the EM morphology of a given cell type. After reconstructing neurons in FAFB we registered these to a common template brain, (JRC2018F, Bogovic et al., 2020) via the natverse::xform_brain function in R, and wrote an image stack of this registered neuron (see https://github.com/jefferislab/skeleton-to-MIP). To compare this stack with existing images of driver line libraries we used the Color depth MIP mask search ImageJ plugin; first to generate a color-coded 2D intensity projection of the stack, and then to compare this with the MIP images of large driver line libraries from the Janelia FlyLight team (Jenett et al., 2012; Meissner et al., 2020; Otsuna et al., 2018; Tirian and Dickson, 2017). We then selected split-GAL4 hemidriver lines labeling our neuron of interest based on the full expression pattern of GAL4 using the same enhancer, and multi-color flip-out (MCFO) labeling of these drivers. Our split-GAL4 lines contain two hemidrivers, the p65ADZp in attP40 and the ZpGAL4DBD in attP2, with a few exceptions where a hemidriver of a non-GMR enhancer was used (JK1029-AD, or *dsx*-DBD). The selected GAL4 and split-GAL4 line candidates were screened via confocal microscopy by combining the two hemidrivers and a UAS reporter: Enhancer-p65ADZp (attP40); Enhancer-ZpGAL4DBD (attP2) crossed to 20xUAS-CsChrimson::mVenus (attP18) or UAS-CD8::GFP; UAS-CD8::GFP.

### Neuron tracing in FAFB

We used a serial section transmission EM volume to sparsely reconstruct the morphology and connectivity of neurons of interest in a female fly brain volume (FAFB, Zheng et al., 2018). Neurons were reconstructed in three ways: 1) fully manual reconstruction (lPN, lvPN, aSP-g): tracing and segment concatenation was done using CATMAID (Saalfeld et al., 2009), a Web-based environment for working on large image datasets and for tracing of neuronal morphologies. Annotated synapses represent chemical synapses based on previously described criteria. 2) To sample the presynaptic partners of aSP-g neurons we used an automated segmentation of the FAFB dataset with manually annotated presynaptic locations (Li et al., 2020). The presynaptic locations were mapped onto the volumetric neuron segments, that allowed us to rank upstream segments by the number of presynapses inside the volume. We traced all upstream segments with more than one presynapse, thereby covering 56% of all inputs to aSP-g neurons. To reconstruct upstream neuron morphologies we concatenated skeletonized versions of the segments as described in Bates et al. 2020. 3) To sample the presynaptic partners of G2N-SLP1 neurons we relied on another automated segmentation of the FAFB dataset, and the related FlyWire proofreading environment (Dorkenwald et al., 2020). We ranked upstream segments by the number of manually marked synaptic locations inside their volume, and all segments containing more than two synapses (75% of all G2N-SLP1 inputs) were reconstructed via merging segments. pC1 reconstructions in FAFB were made publicly available in Wang et al. 2020.

### Computational neuroanatomy and connectomic analysis

A dense reconstruction of one third of a female fly brain imaged with FIBSEM (focused ion-beam scanning electron microscopy), referred to as the hemibrain, was used to investigate connectivity in the antennal lobe: for ORNs, il3LN6, PNs (Scheffer et al., 2020). The website displaying the data (neuprint.org) and the natverse R package family (natverse.org) was used to query connectivity information, and to visualize neuron morphologies (Bates et al., 2020b). Neuron identifiers and the number of synaptic connections across cell types from both datasets can be found in Supplementary File 2. Our group identified neurons in the hemibrain prior to publication and contributed the annotation of all ORNs, PNs, and LH neurons in neuprint (Schlegel et al., 2021).

To count the number of branches in axon cross sections in il3LN6 (Figure 4K) we used the neuroglancer environment of the respective EM dataset (hemibrain or FAFB-FlyWire), and navigated to the EM section where il3LN6 neurites enter the AL. il3LN6 neurons were previously identified in the hemibrain datasets, and we found the corresponding FAFB neurons based on their morphology. The two il3LN6 from the hemibrain provide two data points on Figure 4K, and the two il3LN6 in FAFB provide four data points (two neurons, two hemispheres). We used a one-sample t-test to test whether the mean number of branches is different from 1–the usual number of branches in a neurite that connects distant parts of fly neurons.

To find third-order neurons downstream of lPNs and lvPNs (Figure 6A) we queried the hemibrain:v1.2.1 dataset displayed at neuprint.janelia.org via the neuprintr R package. We selected downstream cell types for further characterization based on a sliding threshold combining the absolute number and the relative fraction of inputs from a given PN type. Cell types with not more than 10 inputs were excluded, and cell types with more than 50 inputs were included, irrespective of their relative PN input. Cell types between 11 and 50 inputs were included if they were above a slope defined by the following points along the absolute and relative input dimensions: 10 synapses, 4% relative input; 50 synapses, 0.5% relative input. For lPNs this analysis was limited to the LH (thereby excluding cell types that get input from dendritic boutons in the AL, and cell types postsynaptic to lPNs in the mushroom body calyx); for lvPNs this analysis was limited to downstream partners in the LH and SIP. To assign projection pattern based classes (LN, ON, DN), and neurotransmitters to these cell types we partly used previous work from our group (Bates et al., 2020a; Dolan et al., 2019; Schlegel et al., 2021). Cell types that were not included in these previous analyses were inspected manually to assign them into projection groups, and a machine learning algorithm was used to predict neurotransmitters based on the ultrastructure of synaptic terminals (Eckstein et al., 2020). To classify the third-order cell types by input selectivity we manually inspected the presynaptic neuron pool of each cell type. We inspected presynaptic cell types that provide either more than 0.5% of synaptic inputs or more than 10 synapses to the respective third-order cell type. These presynaptic neurons were sorted into two groups: sensory and higher-order based on their projections. Neurons with dendrites in the SEZ, anterior ventrolateral protocerebrum, or the optic lobes were classified as sensory, as these neuropils are known to relay gustatory, auditory and mechanosensory, and visual information, respectively. If for a given third-order neuron the sensory input from these pathways was more than 25% of its olfactory inputs from uniglomerular PNs we classified it as ‘multimodal’. The remaining third-order cell types were classified as ‘DA1 selective’ if DA1 PNs provided more than 50% of their olfactory inputs, and ‘mixed olfactory’ if the DA1 PN input was less than that.

For morphological clustering (Figure S7C) we calculated mean NBLAST similarity scores of neuron skeletons (point and line representations) and used Ward’s hierarchical clustering on these scores and expert inspection to find morphological cell types (Costa et al., 2016).

Quantification of dendritic cable in the lateral horn (Figure S7B) was done with the nat R package (natverse.org/nat). Neuron skeletons were resampled at 1 μm, to get an even distribution of nodes throughout the neuronal cable. We pruned these skeletons to dendrites by manually selecting a node on the skeleton before the axon branching, and removing all nodes distal to that. After this we took the number of nodes that were inside the lateral horn, divided by the number of all nodes. For FAFB neurons, we used the LH_L volume (lateral horn left) to define which synapses are inside or outside the LH. For FlyCircuit neurons we used the LH volume of the FCWB reference brain, which is the template that these neurons were registered to in the dataset. For MCFO data, neuron skeletons were traced in Fiji (Schindelin et al., 2012) with the Simple Neurite Tracer plugin (Longair et al., 2011) and then registered to the IS2 template brain with CMTK–Computational Morphometry Toolkit as described in Cachero et al. (2010). We used the LH volume of the IS2 template brain to calculate the dendritic cable inside the LH for neurons from MCFO data.

### Immunohistochemistry and confocal microscopy

Immunohistochemistry was done as described in (Jefferis et al., 2007) except that the blocking step was overnight at 4°C. Primary antibodies: mouse anti-nc82 (DSHB, AB_2314866) 1:40, chicken anti-GFP (Abcam, ab13970) 1:1000, mouse anti-ChAT4B (DSHB, AB_528122), rabbit antiGABA (Sigma, A2052). Secondary antibodies: Alexa-568 anti-mouse (Invitrogen) 1:400, Alexa-488 anti-chicken (Invitrogen) 1:400, Alexa-633 anti-mouse (Invitrogen) 1:400, Alexa-568 anti-rabbit (Invitrogen) 1:400.

Prolonged incubation (2—3 days at 4°C) with primary and secondary antibodies was required for homogeneous staining. Specimens were whole mounted in Vectashield (Vector Labs) on charged slides to avoid movement. Confocal stacks were acquired using a Zeiss 780 confocal microscope. Brains were imaged at 768 × 768 pixel resolution every 1 μm (0.46 × 0.46 × 1 μm) using an EC Plan-Neofluar 40×/1.30 oil objective and 0.6 zoom factor. All images were acquired at 16-bit color depth. Maximum projections of z stacks were made in Fiji (Schindelin et al., 2012).

### In vivo calcium imaging and stimulus presentation

Functional imaging experiments of neurons were performed on virgin female or male flies aged 3 to 7 days, containing one copy of codon optimized GCaMP6f, unless other GCaMP is specified. Flies were placed into custom built holders, leaving the head and thorax exposed, under ice anesthesia and secured in place with UV curable glue (Norland Optical Adhesive, NOA 68). Low melting point wax was used for immobilizing the legs and the proboscis. A window was then cut into the head capsule with a 30G needle, and trachea and air sacks were removed with forceps. Fly brains were bathed in external saline adjusted to 275 mM and 7.3 pH, and bubbled with 5% CO2 - 95% O2 mixture. The saline had the following composition (Concentration, mM): NaCl 104.75; KCl 5; NaH_2_PO_4_ 1; MgCl_2_.6H_2_O 1; CaCl_2_.2H_2_O 1; NaHCO_3_ 26; TES 5; glucose 10; trehalose 10. The antennae were left under the holder so that they could be exposed to odor stimuli. A custom-built setup based on the Sutter (Novato, CA) Movable Objective Microscope with a Zeiss W PlanApochromat 20x/1.0 objective was used for the two-photon imaging. A Coherent (Santa Clara, CA) Chameleon Vision Ti-Sapphire provided 900 nm laser excitation, and image acquisition was controlled by Vidrio ScanImage Premium software (Leesburg, VA Pologruto et al., 2003). Image acquisition and stimulus delivery were triggered by a separate computer via Igor Pro software (Wavemetrics, Lake Oswego, OR) running Neuromatic. Images were captured at 7 Hz at 200 × 200 or 140 × 280 pixels, or at 21 Hz, with two bilaterally placed 80 × 80 pixel ROIs.

cVA was delivered via a custom built olfactometer with two odor channels, each equipped with a solenoid valve (SH360T041, Neptune Research). Carrier airflow rate was 600 ml/min and odor channels entered the airstream approximately 3 cm from the fly’s antennae with a flow rate of 200 ml/min, all regulated by separate mass flow controllers (Alicat Scientific Tucson, AZ, MC Series). Clean air from both odor channels was constantly flowing to the fly until a trigger arrived to one of the valves, redirecting the odorized air from waste to the fly. Odors were 10% cVA (Pherobank, CAS: 6186-98-7, product number: 10421) diluted in mineral oil, and the solvent control. The odor path containing cVA had a manual valve between the mass flow controller and the odor bottle that was used to send the air to waste in between presentations to avoid depletion of cVA from the bottle with constant airflow.

Males used as stimuli were 4-8 days old Canton S flies, collected upon hatching and raised in groups of 5-10 individuals. A single male was selected and had its legs and wings removed under ice anesthesia, and glued onto a metal needle with UV-curable glue (Norland Optical Adhesive, NOA 68). The glue was applied onto the proboscis, thorax, and abdomen of the male to inhibit any movement, but the genitalia were left free to avoid covering the regions where cVA is most abundant. When presenting female flies as a stimulus the same procedure was used with 4-8 days old Canton S virgins, reared in groups. For stimulus calibration a female fly was placed in the imaging holder (and later discarded) to position the male relative to the imaged fly’s antennae. A Mini 23 Luigs Neumann micromanipulator was used to move the male, controlled by an SM-5 system from the same manufacturer. The SM-5 was connected to the imaging PC to externally trigger movements of the stimulus fly to defined locations with custom MATLAB scripts. The male, facing up with its genitalia, was positioned manually directly in front of the female’s antennae by the help of a camera equipped with a high magnification lens (FLIR BlackFly S3, and 3.3X Macro Zoom Lens, Computar). The manipulator was zeroed in this position, so that any subsequent movement of the male happened relative to this origin. Male movement via the manipulator and two-photon imaging was triggered as described above. To infer the timing of male movement a camera (same as above) was triggered together with the imaging experiment, and recorded throughout the acquisition at 33 frames per second. The start and end frames for each movement were noted down, and male movement traces were generated based on these time points in R, assuming constant velocity. An IR LED was used for illumination during imaging, and the camera was protected from 2-photon light with an 800 nm short pass filter.

For all stimulus protocols the starting position of the male was 10 mm below the female’s antennae. For single male presentations the male was moved to 0.75 mm distance for 10 s. For speed tuning experiments, the presentation length at lower speeds was shorter, as the time of movement start (both up and down) was kept constant. We used three speeds: 1.41, 4.30, and 8.04 mm/s, which correspond to speed settings 7, 11, and 15 (maximal) on the micromanipulator, respectively. For distance tuning experiments we used ten distances: 5, 3.5, 3, 2.5, 2, 1.5, 1, 0.75, 0.5, 0.25 mm. For each distance the male was moved up for ~5 s, and then lowered back to the starting positions (10 mm) for ~12 s. For bilateral presentation experiments the male was moved to 0.5 mm distance in z, and 1.25 mm laterally to one side with respect to the antennae. During bilateral presentation responses from both hemispheres were recorded (with the exception of Figure 4M); for ORNs and lPNs in parallel, for lvPNs sequentially. Individual hemispheres were analyzed separately, this resulted in data points twice the number of imaged flies for experiments with intact antennae, and the same number of data points as flies for antennal block conditions. This way blocking an antenna and recording from both sides again results in one hemisphere with its ipsi-, and one with its contralateral antenna blocked. For 2D spatial coding experiments we used sixteen positions defined by a hexagonal lattice centered around the imaged fly (Figure 5A), and recorded responses in both hemispheres in parallel. The points had a distance of 1, 1.732, or 2 mm from the antennae, and an angular position ranging from +150°. We did not use 180° presentations, as the imaged fly’s body takes up these positions defined by the lattice.

Antennae were blocked in the respective bilateral presentation experiments with Kwik-Sil (World Precision Instruments), a fast curing, low toxicity adhesive. The two components of Kwik-Sil were mixed and a small amount of fumed Silica (Sigma, S5130) was added to speed up curing. The mixture was gently applied on one of the antennae under a dissection scope, with care taken not to touch the other antenna. All flies used for these experiments were imaged with intact antennae prior to the antennal block, and the resulting data both pre and post block is included in the relevant figures and analyses (Figure 3, 4).

Optogenetic stimulation of lvPNs via CsChrimson during pC1 imaging (Figure 1 N) was done by a fiber-coupled 617 nm LED (M617F2, Thorlabs, Ely, UK). The light was passed through a 600 nm long-pass filter, to avoid any bleed-through into the imaging PMT (GCaMP emission filter was 525/70 nm band-pass). An optic fiber was placed approximately 0.5 mm away from the fly’s head from below, and the LED was controlled via an external trigger from Igor as described above. The LED stimulated with 50 ms light pulses for 5 s at 10 Hz. To record pC1 activity we imaged a location where only the branches of pC1 neurons are labeled by *dsx-*LexA: the most medial branches in the ROI marked on Figure 6C. To find this location we collected the reconstructions of all *dsx+* neurons in the hemibrain dataset and overlaid them to define a region where pC1 branches are clearly separated and recognizable from the view on the 2P-scope.

Chemogenetic block was performed via expressing the histamine-gated chloride channel, Ort (Figure 4M), under UAS control driven by VT046100-GAL4, a line that labels only il3LN6 neurons in the antennal lobe. The antennal lobe is not innervated by histaminergic neurons therefore Ort can be used as a specific and potent inhibitor of neural activity when expressed in the AL transgenically (Liu and Wilson, 2013). The brain was covered in regular saline while recording control responses. After this the saline was swiftly removed with a Venturi pump, and replaced by pipetting 1 ml of 2 mM histamine-chloride (Sigma H7250-5G) diluted in saline. Responses under histamine block were measured three minutes after histamine application. To wash out histamine, the above procedure was repeated twice with imaging saline, and responses were measured three minutes later. We used VT033066-LexA to drive GCaMP expression in lPNs, and imaged their axons in the ventromedial lateral horn. The most medial part of this area contains almost exclusively DA1 lPN axons, however some other PN types (DL3, VA1v, VA1d) that respond to fly odors and are also labeled by this driver line have arbors in the vicinity. This likely contributed to the more sustained responses observed in these experiments.

### In vivo labellar stimulation and Calcium imaging

Flies used in these experiments were reared on a yeast-based medium as described in Carvalho-Santos et al., (2020). Labellar stimulation experiments (Figure 8D, E) were performed on virgin female flies aged 2 to 7 days, expressing GCaMP7f under the control of Ir94e-GAL4^2^. Flies were fixed to a custom-built acrylic block using UV curable glue (Bondic, Niagara Falls, New York, US). The proboscis was extended using a blunt needle (B30-50; SAI Infusion, Faridabad, Haryana, India) attached to a vacuum pump (N86KN.18; KNF DAC GmbH, Hamburg, Germany) and fixed in an extended position by carefully applying UV curing glue only to the proximal part of the proboscis using an insect pin, such that the labellum could move freely. The front legs were removed to prevent flies from touching the stimulus. The anterior part of the head capsule was placed through a hole in a plastic weigh boat that was fixed on top of the fly. The space between the head and the weigh boat was sealed with UV curable glue. The head capsule was covered with carbogenated (95%O_2_, 5%CO_2_) adult hemolymph-like saline of the following composition (Concentration, mM): NaCl 103; KCl 3; TES 5; trehalose dihydrate 10; glucose 10; sucrose 2; NaHCO_3_ 26; CaCl_2_ dihydrate 2; MgCl_2_ hexahydrate 4; NaH_2_PO_4_ 1; pH 7.3). A window was cut between the eyes and the ocelli, thereby removing the antennae. Trachea covering the brain were removed and the esophagus was transected to allow for unoccluded visual access to the SEZ.

Image acquisition was performed using a resonant-scanning two-photon microscope (Scientifica, UK). The system was equipped with a 20x/1.0 water immersion objective (Olympus, Japan), controlled by a piezo-electric z-focus, allowing for fast volumetric scans. A Chameleon Ultra II Ti:Sapphire laser (Coherent, Santa Clara, CA, USA) was used to excite GCaMP7f at 920 nm. Imaging data were acquired using SciScan (Scientifica, UK). 60 s recordings of the SEZ volume were performed at 1 Hz volume rate covering 512 × 256 × 60 voxels at voxel dimensions of ~0.5 × 0.5 × 3.6 μm. Scanning was performed in sawtooth mode and 5 z-planes acquired during flyback were removed. During imaging, the brain was constantly perfused with saline bubbled with carbogen (95 % O_2_, 5 % CO_2_).

Female and male virgin flies aged 1 to 7 days were glued onto metal needles for stimulations as described above. Water and NaCl (100 mM) stimuli were presented using glass capillaries. Capillaries (GC15F-10, Harvard Apparatus, Edenbridge, Kent, UK) were pulled using a laser pipette puller (P2000; Sutter, Novato, CA, USA) to have a blunt end and an inner diameter fitting the fly proboscis. 200 mL pipette tips were cut to fit the glass capillaries and sealed with Parafilm (Amcor, Zürich, Switzerland). Stimuli were positioned in front of the fly proboscis using a micromanipulator (Sensapex, Finland). Positioning and stimulation were performed under visual control using a PointGrey Flea3 camera and a custom Bonsai script (Lopes et al., 2015). All stimuli were prepared in MilliQ water (Merck KgaA, Darmstadt, Germany). During imaging, two taste stimulations were performed by touching the proboscis with the respective stimulus at 10-15 s and 20-30 s.

### Calcium imaging quantification and statistical analyses

Images were registered in x and y with the NoRMCorre algorithm implemented in MATLAB using the signal channel (Eftychios A. Pnevmatikakis, 2017) https://github.com/flatironinstitute/NoRMCorre. Flies with notable movement in the z axis were removed from analysis. Image analysis was performed with custom scripts written in R employing the open source scanimage package (see https://github.com/jefferis/scanimage). To calculate ΔF/F_0_ we defined F0 as the mean fluorescence value of frames between 1 s after the start of the imaging sweep until the start of the stimulus. ΔF/F_0_ traces were gently smoothed with a low-pass Butterworth filter, except for speed tuning experiments (Figure 7). For distance and angular tuning curves, ΔF/F_0_ values were normalized by the largest value from a given ROI over an experiment. Response maxima and means were calculated in R.

Distance tuning curves were calculated based on the mean normalized response maxima to a given male distance, and fitted with a sigmoid curve. Curve fitting was done with nonlinear least squares method, self-started by a logistic function and parameters from the data in R (Figure 2D, E, F). We compared peak responses to a 10 s presentation across the three imaged fly-presented fly sex pairings (female to male, male to male, female to female) for a given cell type (lPN, lvPN) with Kruskall-Wallis test. This was followed by pairwise Wilcoxon-test with Benjamini-Hochberg correction for multiple comparisons (Figure 2H, I; Figure S2C).

For bilateral presentation experiments (Figure 3, 4) the mean values of normalized traces were taken from six responses for all imaging ROIs. These data were checked for normality (Shapiro-Wilk’s test), and the variance of responses to ipsilateral and contralateral presentations were compared with F-test. If a condition passed both tests (p > 0.05), unpaired t-test was used to test the statistical significance of response differences to ipsi- and contralateral presentations. Where either condition failed (normality, or equal variance), Wilcoxon-test was used instead. To compare bilateral contrast, we took the difference between the mean ipsilateral and the mean contralateral response for a given ROI. To compare differences in bilateral contrast across cell types (Figure 4F) we used Kruskall-Wallis test, followed by pairwise Wilcoxon tests, with Benjamini-Hochberg correction for multiple comparisons. To compare bilateral contrast before, during, and after blocking il3LN6 we used Friedman-test, followed by pairwise paired Wilcoxon tests, with Benjamini-Hochberg correction for multiple comparisons (Figure 4N).

For 2D positional coding experiments we used a hexagonal lattice centered around the imaged fly to define male positions, thereby sampling 2D space at equal distances between neighboring stimulus positions. This resulted in three possible distances from the imaged fly’s antennae: 1, 1.732, and 2 mm, and eleven angular positions (30° steps between −150° to +150° with 0° being frontal to the imaged fly). To create the angular tuning curves of left and right lPNs we used a fixed distance (1.732 mm), for which mean responses at six angular positions were recorded directly, and mean responses at five positions were linearly interpolated based on mean responses to 1 mm and 2 mm presentations at these angles. A single multivariate linear model was used to predict the x and y position (equal to the cosine and sine of the angular position on a unit circle, respectively) of the male based on the difference and the sum of right and left lPN responses. Based on these predictions of x and y position the angular position was calculated and compared with the actual angular position to get prediction errors in degrees and mm (Figure 5G, Figure S5A, B).

For speed dependence experiments, mean response maxima from six trials per fly were compared by Friedman-test to assess if male approach speed had a significant effect on responses. Where the Friedman-test rejected the null-hypothesis (AV2a2) it was followed by paired Wilcoxon-test for pairwise comparisons across speeds with Benjamini-Hochberg correction for multiple comparisons (Figure 7). All analyses were done in R.

Representative 2-photon images with inverted grayscale pixels were made in Fiji (Figure S2B, Figure4G; Schindelin et al., 2012). Calcium imaging data of labellar stimulations were motion corrected using 3dvolreg from the afni toolkit (Cox, 1996). Volumes were then filtered using a 3×3×3px gaussian filter, and collapsed to 2D by performing a maximum intensity projection in python. Using Fiji, circular ROIs were manually drawn around the four Ir94e projection areas in the SEZ, and average time-series information was extracted. ΔF/F0 was then calculated in R and the data was normalized to the maximum value within a fly. Mean values were calculated by averaging ΔF/F0 during stimulation (10-30 s). Stimulus elicited responses were compared using Tukey’s honest significance test.

### Courtship assay, aggression, and behavioral analysis

An assay modified from Hoopfer et al. (2015) was used to measure male courtship, female receptivity, male-male aggression and female-female aggression. For courtship and receptivity experiments, 4-8 day old virgin flies of the experimental genotype, raised in groups of 20 same-sex flies, and 4-8 day old virgin Canton S partners of the opposite sex were placed with gentle aspiration in a transparent behavioral plate with eight chambers, 16 mm in diameter x 12 mm height, equipped with sliding separators. For aggression experiments, two experimental virgin males or females were taken from separate vials. Walls were covered with teflonlike material (polytetrafluorethylene, Sigma-Aldrich 665800-100ml) and the lid was covered with Sigma-coat (Sigma-Aldrich SL2-100ml) to prevent flies from climbing and holding onto the walls and lid. The plate was placed into a 23°C incubator and males and females were allowed to habituate to the chamber for a few minutes after transfer. The separators were removed upon the start of the experiment, and flies behaved and interacted freely in the chambers. The behavioral plate was backlit with homogenous IR light from an LED panel (850 nm), and a FLIR Grasshopper 3 camera was used to record behavior for 20 minutes at 30 frames per second. For some experiments, bright or dim ambient light was provided to the flies to stimulate courtship by the males, while other experiments were done in complete darkness (see figures for light conditions: light on, dim, or darkness). The intensity of the ambient light was adjusted in experiments for a given cell type manipulation for all conditions, to set the baseline level of courtship and copulation. This was necessary to avoid situations where genetic controls mated immediately, in which case a receptivity increase by a manipulation could not be detected due to a ceiling effect.

Video files were converted to a compressed format (micro-Fly Movie Format, ufmf, Robie et al., 2017) and fly positions were tracked with FlyTracker software (Eyjólfsdóttir, 2014). Tracking data was fed into a JAABA analysis pipeline with custom behavioral classifiers, also detecting the time of mating (Kabra et al., 2013). We trained classifiers for mating, wing extension (as proxy for male courtship, Figure 1H, J, M, O), lunges (as proxy for male aggression, Figure 1L, M), and wing threats (as proxy for female aggression, Figure S1D, G, Supplementary Video 2, Figure 8I). Trained classifiers were tested with ground truth data until high accuracy was achieved compared to an expert annotator. Mating was defined when both flies in the chamber were classified as mating for at least 30s, and mating events were eventually manually checked and corrected for errors. Survival analysis of mating latency, followed by logrank test, was used to test statistical significance of differences in latency to copulation. When multiple comparisons were made, it was followed by post-hoc Benjamini-Hochberg corrections. Data processing was done in MATLAB, statistical analyses were done in R with the survminer package (https://rpkgs.datanovia.com/survminer/index.html).

For neuronal manipulations we used driver lines specific to the neuron of interest and expressed an actuator (UAS-CsChrimson) or a pair of apoptosis promoting proteins (UAS-Hid, UAS-Reaper) to genetically ablate neurons. The same LED panel that provided IR light was equipped with 627 nm LEDs as well to activate CsChrimson. The activation LEDs provided light intensity of 8 μW/mm^2^. For most experiments we used pulsed activation: 5 s long periods of 50 ms light pulses at 10 Hz, separated by 5 s no light, throughout behavior. For female aSP-g activation and for male-male PN activation we used constant light for the duration of the recording. Ambient light inside the incubator was either off or on, see figures. Kir2.1 and genetic ablation via Hid and Reaper were constitutively expressed. Genetic controls carried an empty GAL4 insertion (or split-GAL4, where a split-GAL4 line was used) at the same landing site where the driver was inserted (attP2 for GAL4 lines, and attP40 and attP2 for split-GAL4 hemidrivers), crossed to the same UAS or LexAop effector as experimental groups. For cases where a non-GMR or non-VT hemidriver was used (JK1029-AD, Figure 8), the genetic control carried this transgene together with an empty-DBD. See Key Resources Table for fly stocks and Supplementary File 1 for exact genotypes.

### Opposite sex preference and turning behavior (Figures 2, 3)

4-8 old days wildtype (Canton S strain) stimulus flies were anesthetized on ice and decapitated, then waxed onto 16 mm courtship chambers (same as above), 4 mm from one side. Experimental virgin flies were raised in groups of 20, as described above, then individuals were gently aspirated into the chambers, and kept separate from the stationary fly until sliding bars were opened at the start of the recording. Fly behavior in complete darkness was tracked for 20 minutes with Caltech FlyTracker as above, and tracked features including body center coordinates, velocity, and direction were used to analyze fly behavior. To analyze turning behavior, immobilized male centroid coordinates were manually added to the tracker output files in order to compute the facing angle in relation to the male. Data processing, plots and statistical analysis were done with custom scripts in MATLAB, boxplots in Figures 1, 3 and 7 were produced in R. For OSP calculation (Figure 2L-O), heat maps of time spent in each 1 mm2 bin (Figure S2C), and for turning behavior (Figure 3B-G), we excluded frames where the fly was less than 2 mm away from the rims, where tracking is suboptimal due to flies potentially climbing on the walls. For turning analysis (Figure 3), we defined turn initiation (time 0) when angular velocity became greater than 60°/s, with at least 1 s gap between consecutive turns. Turn maximum (time MAX) is the maximal facing angle reached within 3s from time 0. We excluded turns that crossed facing angle 180° (facing away from target) or 0° (facing in front of target) as the facing angle is a symmetrical measure, and does not reflect the actual change in angle in these circumstances. We analyzed the changes in facing angle made either inside or outside cVA detection range of 2-5 mm around the stationary male, excluding turns made within and during 30 s after leaving the 0-2 mm region, to exclude possible effects on turning direction due to tactile communication, or memory of such communication. We also excluded turns in which the change in facing angle was disproportionately greater than the angular velocity. There is usually a high correlation between those parameters, unless the observer fly passed close to the stimulus fly without changing its direction. We defined the cases to remove when the distance between delta facing angle to the correlation line was greater than 3 standard deviations away from the correlation line. We defined turns away from the stationary male as positive, and counted all positive turns to find their proportion. Credit for fly images used for Figure 3A: Copyright Malcolm Storey /www.discoverlife.org, used according to published policy.

## Author Contributions and Notes

Conceptualization, methodology and writing: IT, DSG, and GSXEJ; Investigation: IT, DSG, ED, DM, SNB, WJM, KIM, KSM IV, MG; visualization: PS; Resources: CR, GSXEJ; Supervision: GSXEJ, CR, and DSG; Funding Acquisition: GSXEJ and CR

The authors declare no conflict of interest.

This article contains supporting information online.

## Acknowledgments

We are grateful to K Eichler, S Hampel and A Seeds, and C H Kang and J Kim for their contributions to tracing FlyWire Ir94e GRNs. We thank Y Aso, M Dolan, Y Jung, B Pfeiffer, D Anderson, and C Potter for sharing unpublished fly stocks, and R Wilson, R Benton, J Carlson, B Dickson, R Yang, M Murthy, D Deutsch, T Kiya, and the Bloomington Stock Center for fly stocks; K Asahina, E Hoopfer and D Anderson for advice on behavioral paradigm and providing blueprints for mating chambers; J-C Billeter for sharing protocols; R Wilson, B Hedwig and members of the Fly Module led by G Maimon at the Woods Hole NS&B course for helpful discussions, S Holtz for advice on pharmacology experiments; members of the Jefferis group, K Vogt, R Benton, T Branco, and J Kohl for comments on the manuscript. This work was supported by an ERC Consolidator grant (649111), core support from the UKRI Medical Research Council (MC-U105188491), NeuroNex2 (2014862) and a Wellcome Trust Collaborative Award (203261/Z/16/Z) to GSXEJ; Marie Curie individual (H2020-IF-748478) and EMBO long-term (ALTF 164-2016) fellowships to DSG; EMBO long-term (ALTF 462-2015) and Sir Henry Wellcome Postdoctoral (110232/Z/15/Z) fellowships to ED; “la Caixa” Banking Foundation (LCF/PR/HR17/52150002), FCT – Fundação para a Ciência e Tecnologia (PTDC/MED-NEU/4001/2021) and Fundação Champalimaud for CR and DM.

## Supplemental Information

**Fig. S1.**
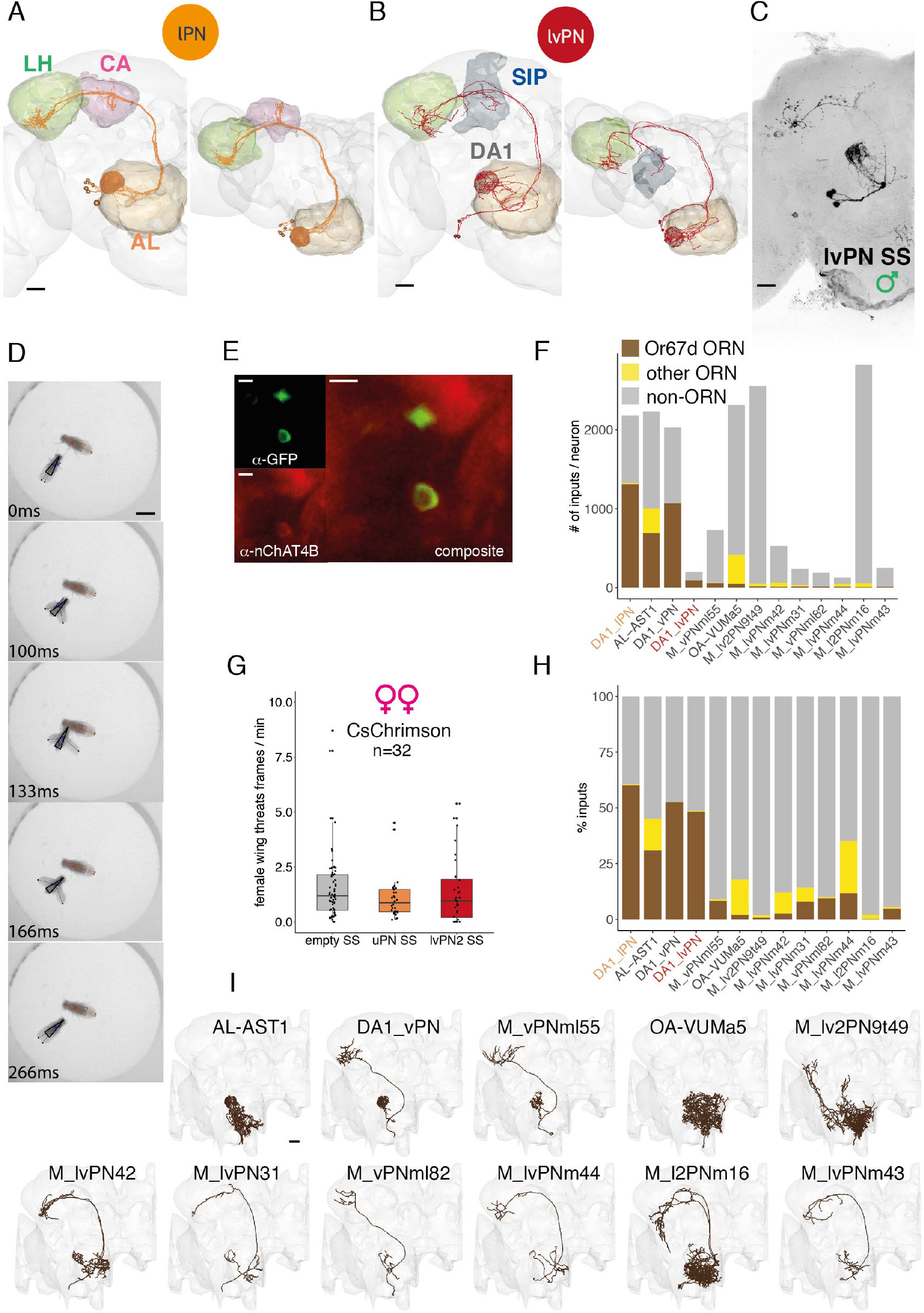
Related to Figure 1. **A, B**: EM reconstructions of DA1 lPNs (A) and DA1 lvPNs (B) in the FAFB dataset. Left: frontal view, right: top view. Scale bars: 20 μm. **C:** Confocal image of DA1 lvPN in a male brain, reporter expression driven by lvPN Stable Split, maximum projection. Scale bar: 20 μm. **D**. An example of female-female behavioral sequence classified as Wing Threat by JAABA classifier. Scale bar: 2 mm. **E:** nChAT4b and lvPN soma co-immunostaining. Top left: lvPN-SS x CD8::GFP - anti-GFP staining. Bottom left: anti-nChAT4B staining, Right: composite image. Scale bars: 5 μm. **F:** The number of inputs per neuron for all non-ORN and non-LN cell types that have more than 10 inputs from DA1 ORNs in the hemibrain dataset. Brown: DA1/Or67d ORN input, yellow: other ORN input, gray: non-ORN input. **G:** Optogenetic activation of lPN or lvPN in pairs of females, using constant red light, 627 nm, 8 μW/mm2 for 20 min. There was no change in female-female aggression. Kruskal-Wallis rank sum test p = 0.27. **H:** The ratio of inputs for the same cell types as in F. Brown: DA1/Or67d ORN input, yellow: other ORN input, gray: non-ORN input. **I:** DA1 ORN downstream cell types ordered by the number of DA1 ORN inputs per cell type, lPNs and lvPNs are not shown. Scale bar: 20 μm.

**Fig. S2.**
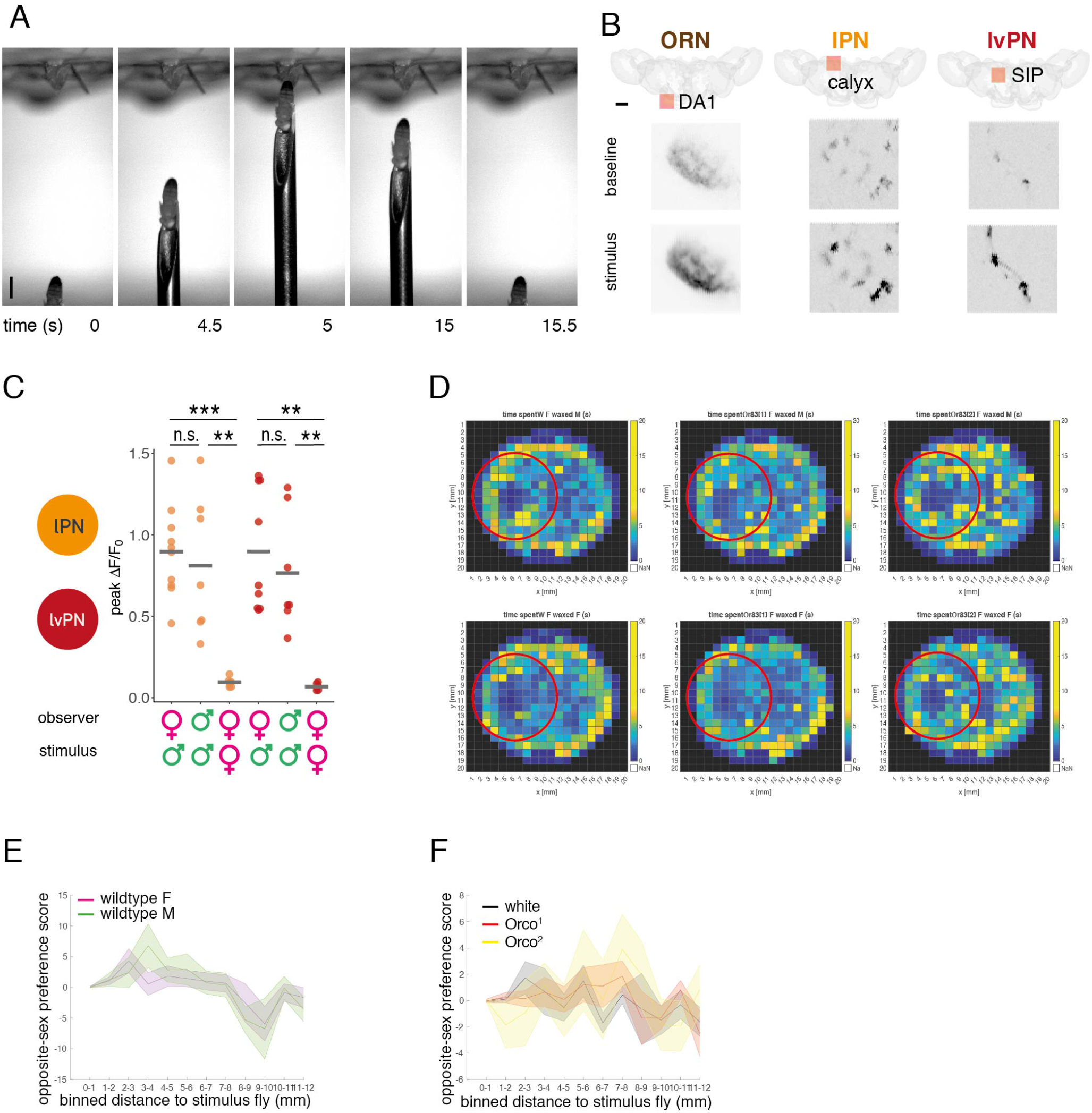
Related to Figure 2. **A**: Image sequence from a video of a single male presentation. A fixed female fly is placed in a holder for two-photon imaging (top). A male fly is glued to a needle that is moved by an externally controlled micromanipulator. Scale bar: 1 mm. **B:** Locations (top row) and example GCaMP fluorescence images (bottom row) of imaging ROIs for ORN, lPN, and lvPN imaging. Fly brains are shown from a top view, which is also the perspective of the imaging objective; orange squares show the location of the ROIs; (ORN: DA1 glomerulus, lPN: calyx, lvPN: SIP). The representative images are averages of frames corresponding to 1 s, before (baseline) and immediately after (stimulus) a male fly was presented at 0.75 mm distance as shown in (A), (also described in Figure 2G). Pixel gray level shows GCaMP signal intensity. Scale bar: 40 μm. C: Quantification of responses in Figure 2H, I. The sex of the imaged and the stimulus fly is indicated under the x axis. Mean peak responses from individual flies are shown as points, means across conditions are shown by horizontal bars. lPN data is in orange, lvPN in red. Peak responses in male and female flies are not significantly different in either PN types, responses to a female fly in females are significantly smaller than responses to a male. D: Heat maps of time spent in each 1 mm2 bin in the arena during 20 min recording of an observer virgin female with an immobilized stimulus. Red circles show the area 5 mm around the stimulus, used to calculate OSP for figure 2N. Top line: wildtype male stimulus, bottom line: wildtype female stimulus. Observer female shown, from the left panel: *white, Orco*^1^, *Orco*^2^. E: OSP at 1mm binned distances from stimulus: for both wildtype males and females, OSP was highest in bins within 5 mm distance from stimulus (2-3 mm for females, 3-4 mm for males). Lines represent mean OSP within discrete 1 mm bins. Shaded area is SEM. F: OSP at 1mm binned distances from stimulus: white females had the highest OSP within 2-3 mm distance from stimulus, similar to wildtype females, whereas *Orco*^1^ and *Orco*^2^ females shifted their OSP to greater distances from a stimulus (highest at 8 mm for both). Lines represent mean OSP within discrete 1 mm bins. Shaded area is SEM.

**Fig. S3.**
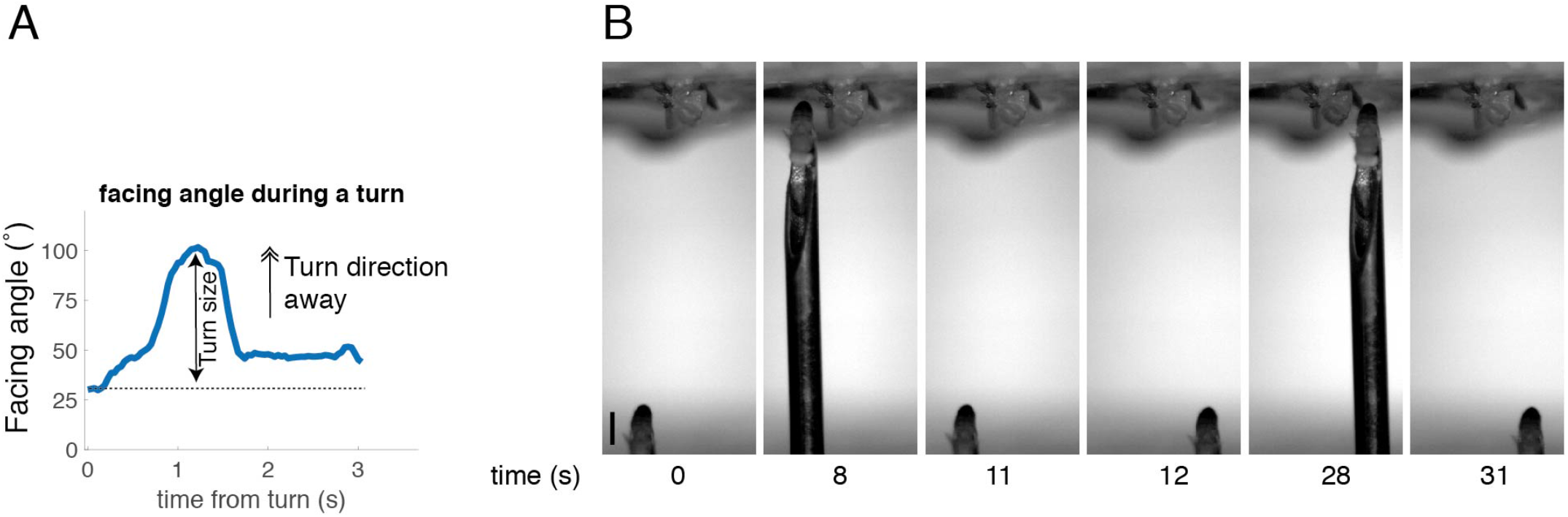
Related to Figure 3. **A:** Facing angle change over time for a single fly during a single turn. Facing angle at time 0 is 30°. Facing angle at time MAX is 101°. Therefore the turn size is 69° and the turn direction is away. **B:** Image sequence from a video of a bilateral male presentation. A fixed female fly is placed in a holder for two-photon imaging (top). A male fly is glued to a needle that is moved by an externally controlled micromanipulator. Scale bar: 1 mm.

**Fig. S4.**
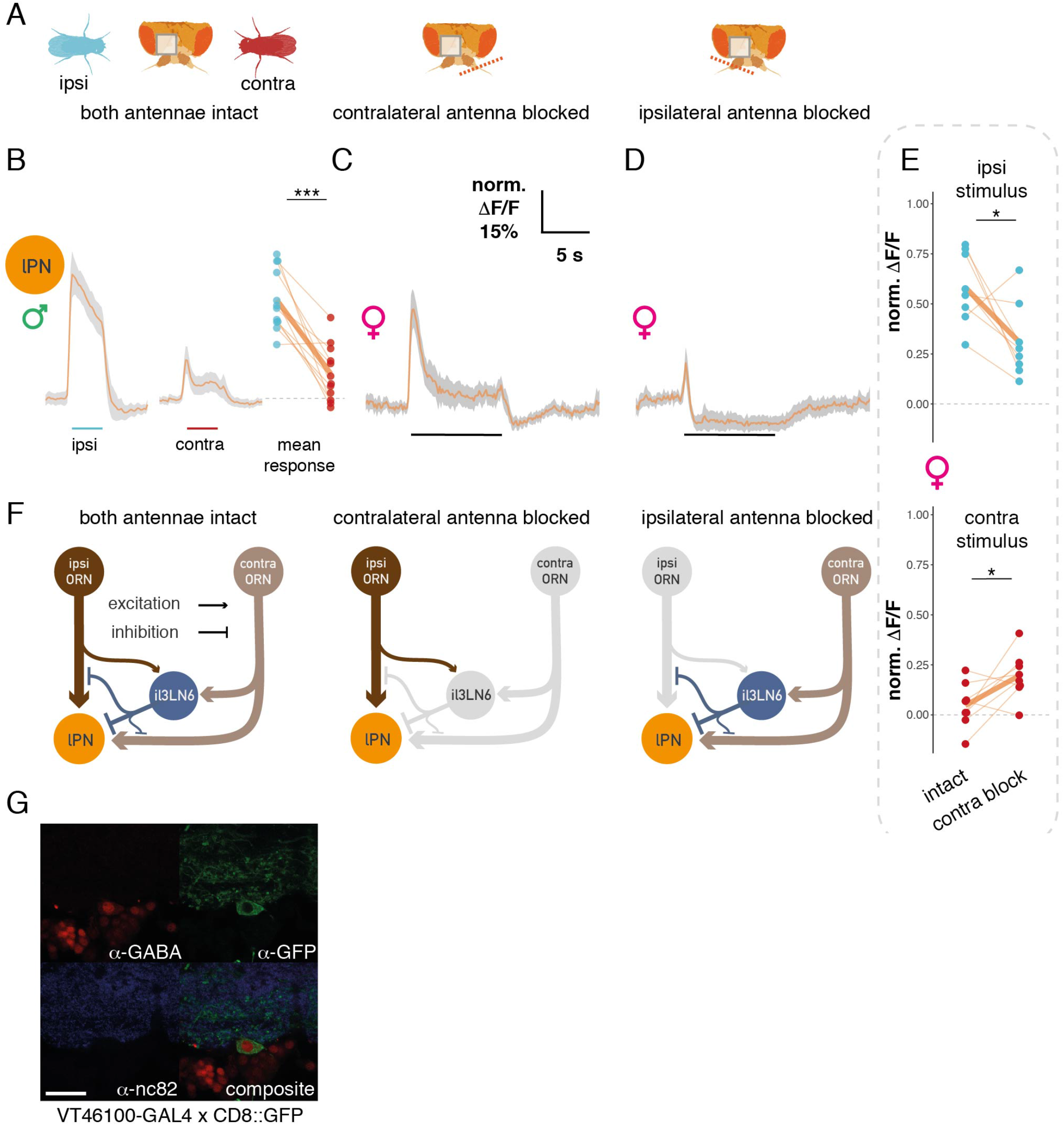
Related to Figure 4. **A:** Schematics of antennal manipulations with respect to an imaging ROI (gray square). Left: both antennae intact, middle: contralateral antenna relative to the imaging ROI is blocked, right: ipsilateral antenna relative to the imaging ROI is blocked. Cyan and red fly cartoons show the relative position of ipsi- and contralateral male presentation, respectively. **B:** Male lPNs show similar responses to female lPNs to bilateral male presentation. GCaMP6f responses in lPN axons to ipsilateral and contralateral male presentation. The male is presented during the time marked by cyan (ipsilateral stimulus) and red (contralateral stimulus) horizontal bars. Mean traces from 6 flies (11 hemispheres), 6 presentations on both sides, gray area is SEM. Left: mean responses of hemispheres during ipsi- and contralateral stimuli **C:** lPN GCaMP6f responses to a male presented in the middle after blocking the contralateral antenna (same stimulus as in Figure 2G). **D:** Same as C, but the ipsilateral antenna is blocked. **E:** Effect of contralateral antenna block on lPN responses to ipsilateral (top) and contralateral (bottom) male presentation. Responses in the same ROIs were compared with paired t-tests before and after blocking the contralateral antenna. Data from Figure 4B, C; n = 8. **F:** Schematic of the expected circuit consequences of antennal block manipulations. Same circuit as in Figure 4O. Left: intact antennae, all circuit elements are functional. Middle: contralateral antenna blocked, contralateral ORNs are not functional, as a result il3LN6 are silent. Ipsilateral antenna blocked: ipsilateral ORNs are not functional, stimulating the contralateral antenna results in parallel excitation and inhibition of lPNs; the net effect of this depends on the position of the stimulus as shown in Figure 4D, see also (D). **G:** GABA and il3LN6 soma co-immunostaining. Top left: a-GABA staining; top right: - VT046100-GAL4 x CD8::GFP, anti-GFP staining; bottom left: anti-nc82 staining; bottom right: composite image. Scale bar: 20 μm.

**Fig. S5.**
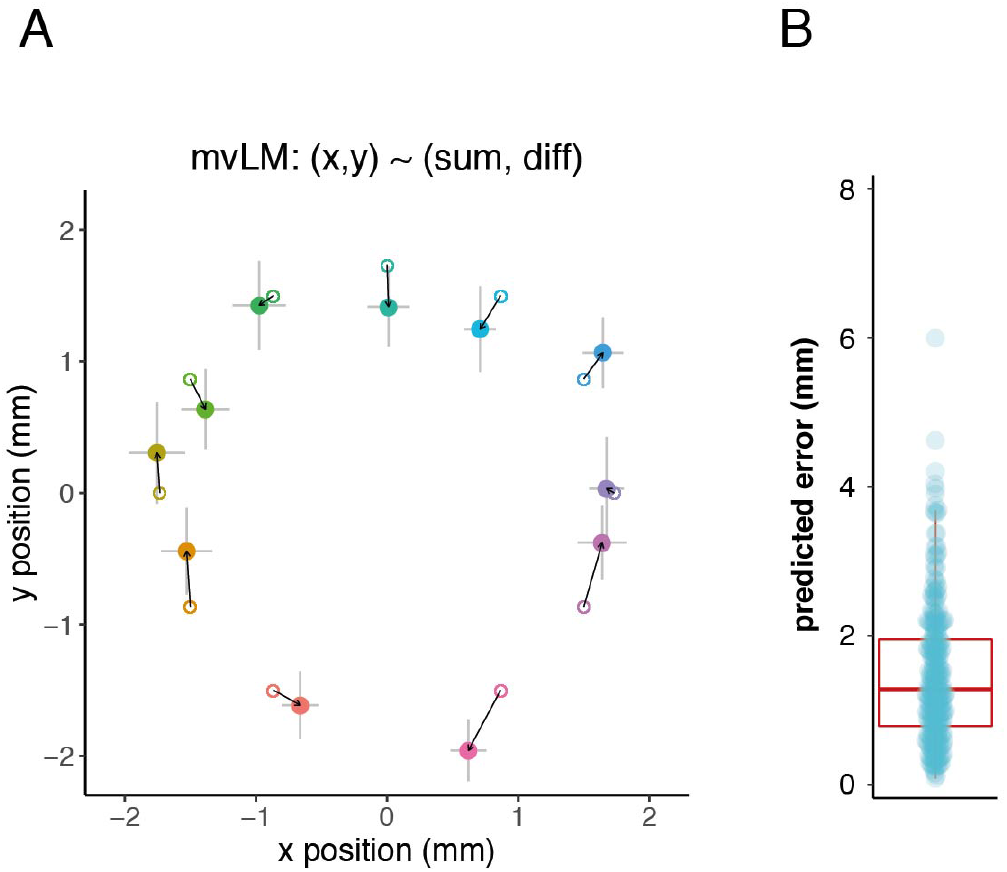
Related to Figure 5. **A:** Male position predictions based on the model described in Figure 5G. Small circles show the original positions; large points show the mean predictions by positions of the model with SEM error bars. mvLM: multivariate linear model; x, y correspond to the x and y coordinates of the male stimulus, with the imaged fly’s antennae as origin; sum and diff correspond to the sum and the difference of right and left lPN responses. **B:** Mean error of position prediction in mm, mean: 1.46, median: 1.28.

**Fig. S6.**
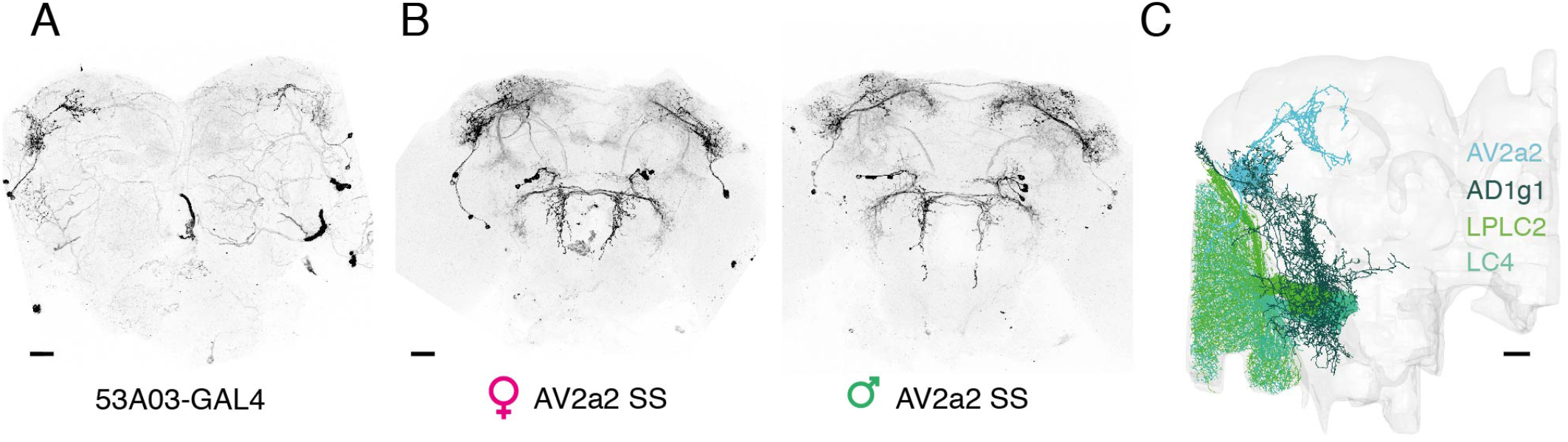
Related to Figure 7. **A:** Confocal image of AV2a2 in a female brain, reporter expression driven by 53A03-GAL4, (used in calcium imaging experiments), maximum projection. Scale bar: 20 μm. **B:** Confocal image of AV2a2 in a female and male brain, reporter expression driven by AV2a2 stable split, (used in behavior experiments), maximum projection. Scale bar: 20 μm. C: EM morphology of AV2a2, AD1g1, LPLC2, and LC4 neurons in the hemibrain dataset. Their schematic connectivity is shown in Figure 7J. Scale bar: 20 μm.

**Fig. S7.**
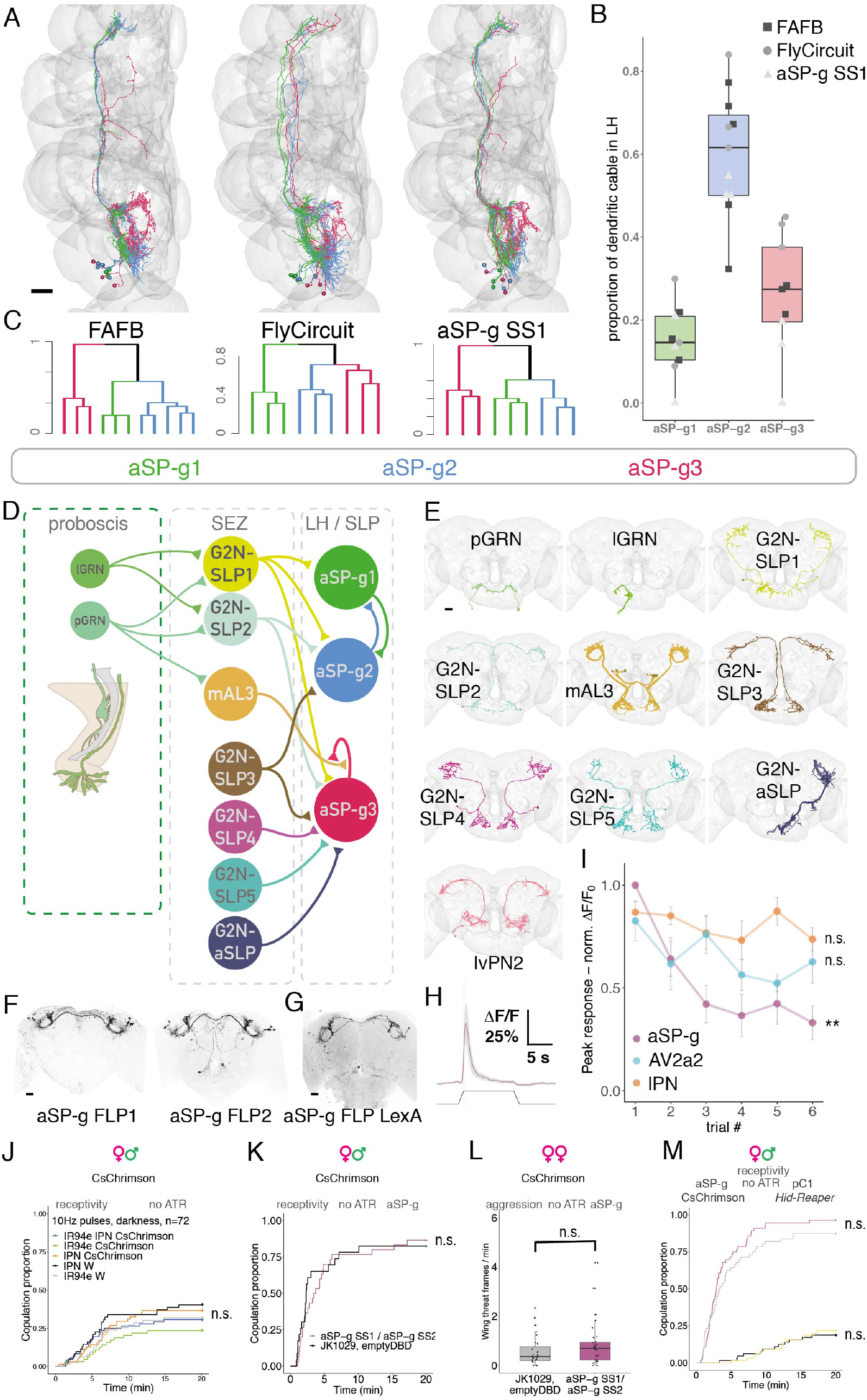
Related to Figure 8. **A**: Reconstructions of aSP-g neurons colored based on NBLAST morphological clustering from the FAFB dataset (left), the FlyCircuit dataset (middle), and MCFO data from aSP-g Stable Split (right). Clusters were named aSP-g1, aSp-g2, and aSP-g3 based on dendritic arbor position from anterior to posterior. Scale bar: 20 μm. **B:** The proportion of dendritic cable inside the lateral horn for aSP-g subtypes across three datasets. Squares: FAFB, circles: FlyCircuit, triangles: aSP-g SS MCFO. **C:** Hierarchical clustering based on NBLAST morphological similarity scores using Ward’s method, k = 3. Order and colors are the same as in A. **D:** Connectivity diagram of aSP-g subtypes with gustatory pathways based on presynaptic sampling of aSP-g dendrites in the FAFB dataset, and reconstruction of G2N-SLP1 inputs via FlyWire. G2N: gustatory second-order neuron, SLP: superior lateral protocerebrum. **E:** EM reconstruction of aSP-g input neurons in the FAFB dataset. First row left: pharyngeal GRNs, first row middle: labellar GRNs, first row right: G2N-SLP1, second row left: G2N-SLP2, second row middle: mAL3, second row right: G2N-SLP3, third row left: G2N-SLP4, third row middle: G2N-SLP5, third row right: G2N-ascendingSLP, Fourth row: lvPN2 (cell types M_lvPNm42 and M_lvPNm44 in the hemibrain dataset); in FAFB, lPN and lvPN2 provide 6.4% and 5.6% of aSP-g2 inputs, respectively. See Supplementary Spreadsheet 2 for synaptic weights for all other connections. Scale bar: 20 μm. **F:** Confocal image of aSP-g FLP1 and aSP-g FLP2 lines in a female brain, (see table S1, genotypes), used to block aSP-g neurons by expressing Kir2.1 (Figure 8H) maximum projection. Scale bar: 20 μm. **G:** Confocal image of aSP-g FLP LexA in a female brain, (see table S1, genotypes), used to activate aSP-g while ablating pC1 for neuronal epistasis (Figure 8K), maximum projection. It is important to note that aSP-g FLP-LexA weakly labels a few neurons in the midline of the brain, projecting from the peri-esophageal region to the pars intercerebralis (although the labeling of these neurons is not strong enough to appear on a maximum projection of the full brain). We cannot exclude the possibility that these neurons contribute to the behavioral effects observed in the behavioral epistasis experiment in Figure 8K. Scale bar: 20 μm. **H:** GCaMP6f responses in aSP-g axons to presenting a male at 0.75 mm distance from the antennae, female fly imaged. Mean trace from 6 flies, 6 male presentations each, gray area is SEM. Same stimulus as in Figure 2G. **I:** aSP-g responses adapt over time, unlike IPN and AV2a2. Normalized peak GCaMP6f responses to a single male presentation (as described in Figure 2G) in 6 consecutive trials separated by 35 s in three cell types, aSP-g: purple, lPN: orange, AV2a2: cyan. Responses are significantly different in aSP-g in different trials (Friedman test), but not in lPN and AV2a2. The same lPN and AV2a2 data were also used in Figure 2H and Figure 7E, F. **J:** control for Figure 8F, using the same experimental conditions but these flies were not raised on all-trans Retinal (ATR) containing food. n = 72 per group. **K:** control for Figure 8G, using the same experimental conditions but these flies were not raised on ATR containing food. n = 32 per group. **L:** control for Figure 8I, using the same experimental conditions but these flies were not raised on ATR containing food. n = 32 per group. **M:** control for Figure 8K, using the same experimental conditions but these flies were not raised on ATR containing food. Notice that pC1 ablation impaired female receptivity regardless of ATR as the ablation is not temporally controlled. n = 64. Throughout the figure, n.s. p>0.05.

## Supplementary File 1

Sheet 1: Statistical tests and significance measures. Where more than two conditions were compared the pairwise comparisons are indicated in the third column; in these cases, Benjamini-Hochberg adjustment was used to correct for multiple comparisons. The number of flies for behavioral experiments represent the group size per group, unless the size differs between groups, in these cases the group size for each genotype is shown separately.

Sheet 2: Full genotypes of flies used in experiments. See Key Resources Table for the genotypes of fly stocks.

## Supplementary File 2

Sheet 1: Connectomics identifiers. In FAFB these are called skeleton IDs and they can be used to look up neuron morphology and connectivity, hosted by Virtual Fly Brain at https://catmaid-fafb.virtualflybrain.org/.

Sheet 2: Connectivity in FAFB and hemibrain datasets. Values indicate the number of synaptic connections between two cell-types in a specific direction; column *from* indicates the presynaptic, and column *to* indicates the postsynaptic cell type. FAFB connectivity is based on partial reconstructions except for the cases where both pre- and postsynaptic arbors were traced to completion. For bilateral neurons the sum of left and right number of synapses in FAFB may not add up to the total connectivity because they reflect connectivity between neurons from the same side, whereas bilateral neurons may connect to neurons from both sides. Hemibrain connectivity reflects version 1.2.1 of the dataset, as displayed at https://neuprint.janelia.org/.

Sheet 3: Third-layer cell types related to Figure 6. Column names indicate the following: *from:*presynaptic PN type; *to*: postsynaptic third-order cell type; *weight:* number of synaptic connections from PN type to third-order cell type; *percent_input:* the percentage that PN inputs make up of all inputs to the given third-order cell type; *projection class:*categories based on where the third-order cell type projects (LN: local neuron, ON: output neuron, DN: descending neuron); *DA1_input_ratio:* the fraction of olfactory inputs that are made by the DA1 PN type; *DA1_selective:* categories based on the DA1 input ratio, if over 0.5, the cell type is DA1 selective, else it is mixed olfactory; *multimodal:* describes whether the amount of non-olfactory sensory input to this cell type is more than 25% of all olfactory inputs; *bilateral*: logical; *NT*: main neurotransmitter, see also methods. *Common_downstream:* logical, if the third-order cell type is downstream to both lPNs and lvPNs; *final_class:* our class of the cell type that uses information from columns *projection_class, DA1_selective, multimodal*.

## Supplementary Video 1

3D rendering of cell types in the first three layers of the cVA circuit in the hemibrain dataset.

## Supplementary Video 2

Virgin female-female aggression during constant aSP-g activation, related to Figure 8I.

